# Hippocampal stimulation timed to memory reactivation shapes human sleep oscillatory dynamics and consolidation

**DOI:** 10.64898/2026.09.02.748628

**Authors:** Prince Okyere, Junheng Li, Tobias Raufeisen, Ketevan Alania, Daniella Louise Jones, Melanie Steiner, Esra Neufeld, Niels Kuster, Nir Grossman, Derk-Jan Dijk, Roi Cohen Kadosh, Ullrich Bartsch, Valeria Jaramillo, Ines R. Violante

**Affiliations:** School of Psychology, University of Surrey, Guildford, UK; School of Biomedical Engineering and Imaging Sciences, King’s College London, London, UK; Department of Brain Sciences, Imperial College London, London, UK; UK DRI at Imperial College London, London, UK; Surrey Sleep Research Centre (SSRC), University of Surrey, Guildford, UK; Defitech Chair of Clinical Neuroengineering, Neuro-X Institute (INX), École Polytechnique Fédérale de Lausanne (EPFL), 1202 Geneva, Switzerland; Foundation for Research on Information Technologies in Society (IT’IS), Zurich, Switzerland; Department of Information Technology and Electrical Engineering, Swiss Federal Institute of Technology (ETH), Zurich, Switzerland; UK DRI, Care Research & Technology at Imperial College London and University of Surrey, UK

## Abstract

Memory consolidation during sleep depends on the precisely timed coordination of hippocampal reactivation with cortical slow oscillations and spindles. In humans this coupling has been characterised correlationally, while causal investigations have required intracranial stimulation in restricted patient cohorts or used non-invasive approaches targeting cortical regions, often obscuring the underlying sleep rhythms. Here we stimulated the human hippocampus during sleep for the first time, applying temporal interference stimulation to the left hippocampal head either concurrently with or prior to induced memory reactivation. Stimulation concurrent with reactivation reduced associative memory forgetting relative to mistimed stimulation and enhanced fast spindle amplitude, preserving the association between slow oscillation–spindle coupling and memory. This behavioural benefit was mediated by a distributed, left-lateralised set of spindle and coupling features. Non-invasively engaging deep hippocampal circuits during sleep offers both a means to probe human memory and a scalable basis for intervention where consolidation fails.

## INTRODUCTION

Whether and how memories endure depends on a process known as consolidation. The active systems consolidation hypothesis offers a framework through which newly encoded hippocampal memories are gradually transferred to stable neocortical representations during non-rapid eye movement (NREM) sleep^1–4^. This is thought to be instantiated by specific electrophysiological signatures: hippocampal reactivation during sharp-wave ripples (80–120 Hz in humans)^5–7^, thalamocortical sleep spindles (9–16 Hz) facilitating information transfer, and cortical slow oscillations (SOs, 0.5 – 2 Hz) organising the temporal structure within which these events unfold^3,8^. Manipulation of hippocampal ripples in rodents demonstrated that memory is impaired if ripples are blocked^9^ and enhanced when prolonged^10^. Yet evidence for similar mechanisms in humans remains correlational. The precise temporal coordination of these rhythms during sleep also supports memory consolidation. Stronger cortico-thalamo-hippocampal coupling, whereby ripples occur during spindles (particularly fast spindles, 12– 16 Hz) nested within SO up-states, is thought to create optimal conditions for synaptic plasticity, and predicts better memory retention^6,11–13^. In rodents, thalamic or cortical stimulation timed to hippocampal ripples or thalamic spindles engages this hierarchy and enhances memory^14–16^. In humans, prefrontal deep brain stimulation synchronised to medial temporal lobe SOs improved ripple-thalamocortical coupling and recognition memory compared with mistimed stimulation^17^.

While valuable, these studies have inherent limitations. Rodent sleep architecture differs from humans, and human intracranial studies typically rely on patients with pharmaco-resistant epilepsy, where small samples, cognitive comorbidities, and medication effects on sleep continuity and architecture complicate interpretation. Non-invasive access to deep brain circuits would transform our ability to causally probe sleep-dependent consolidation mechanisms in the healthy human brain and lay the groundwork for clinically scalable memory interventions. In this context, temporal interference stimulation (TIS) represents a promising approach for selectively targeting deep structures^18^, with recent studies demonstrating it can modulate hippocampal activity during waking tasks and influence memory performance^19,20^.

Here, we tested whether TIS delivered to the human hippocampus in conjunction with induced memory reactivation during sleep, impacted consolidation and the electrophysiological signatures through which sleep supports memory. Across behaviour and sleep electrophysiology we find converging evidence that TIS applied concurrently with induced reactivation reduced post-sleep forgetting compared with stimulation that preceded reactivation. This effect was accompanied by increased fast spindle amplitude, stronger SO-spindle coupling, with the behavioural benefit partly reflected in a coordinated set of sleep EEG features lateralised to the stimulated left hemisphere.

## RESULTS

### Hippocampal TIS during sleep preserves EEG and sleep architecture

Twenty-eight participants (mean age 24.8 ± 6.0 years, 15 males) completed an episodic memory task before and after an afternoon nap (**Fig. 1a**). To induce memory reactivation, we used targeted memory reactivation (TMR), in which cues associated with learned items are re-presented during NREM sleep, a procedure shown to evoke hippocampal activity, reinstate memory-related neural activity, and selectively improve retention of cued over non-cued items^21–24^. Pre-nap, participants learned 120 unique image-word associations, followed by a recognition test in which all studied words were presented intermixed with 40 novel words. For each word, participants made an old or new judgement, yielding a measure of word recognition memory. For items recognised as old, they identified the associated image’s category, face or object, rated their confidence, and described the image, yielding measures of associative recall and memory fidelity respectively. Only items correctly recalled pre-nap were retained and divided equally into four within-subject conditions **(Fig.1b)**: (1) TIS+Cue, where 4 seconds of 90 Hz TIS was delivered simultaneously with the auditory cue; (2) TIS+Cue delayed, where 90 Hz TIS preceded the cue, so that the envelope no longer coincided with reactivation; (3) Cue only, with no TIS; and (4) No-Cue, with no auditory reactivation during sleep and therefore no corresponding cue-locked EEG, serving as the non-cued control condition. All cued conditions were delivered during stable N2 and N3 sleep in pseudo-randomised block order (**Fig. 1c**).

**Figure 1.**
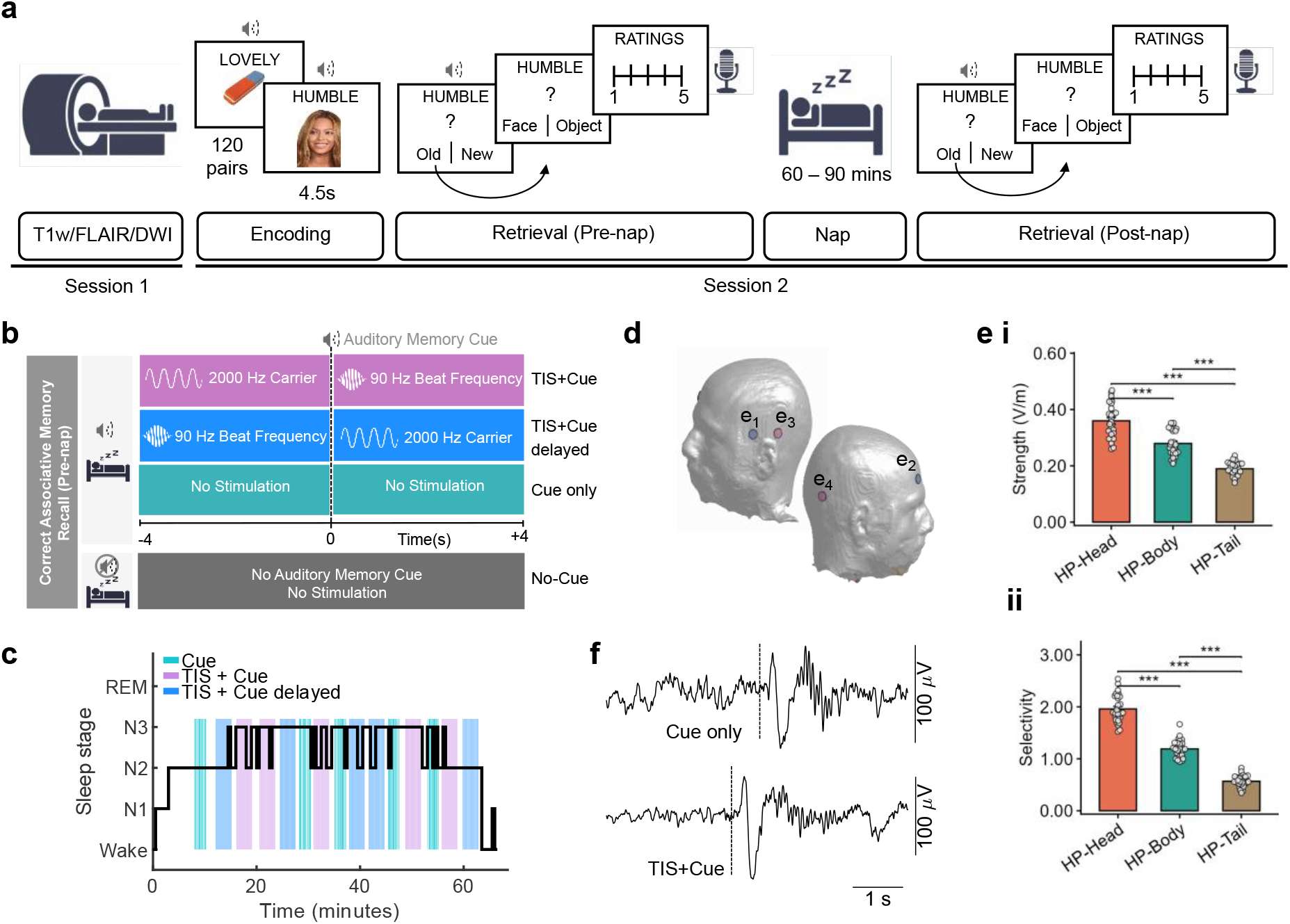
Experimental design and hippocampal TI field metrics: **a.** Experimental design. Each participant took part in two sessions, MRI acquisition (T1w, T2w-FLAIR and Diffusion Weighted Imaging (DWI)) in session 1 and sleep in session 2. In session 2, participants were presented 120 word-image paired associates (each for 4.5 s). Images were faces of famous people or objects, words were adjectives, presented as text and audio (for example, “Lovely” paired with an “eraser”). During the subsequent pre-nap test, participants heard each adjective and judged whether it was paired with the old (studied) or new (novel) stimulus, identified the associated category (face or object), rated their confidence (1–5), and spoke to describe the chosen image. Participants then had a 90-minute afternoon nap opportunity during which temporal interference stimulation (TIS) and targeted memory reactivation (TMR) were applied. Upon waking, participants performed a post-nap test to assess memory retention. **b.** Experimental conditions. Only word-image pairs correctly recalled during pre-nap retrieval were assigned to four conditions (equally distributed). In the TIS+Cue, TIS+Cue delayed, and Cue only conditions, an auditory cue was delivered during the nap time-locked to auditory memory cue onset (t = 0). In the TIS+Cue condition, the 90 Hz beat frequency was delivered simultaneously with auditory memory cue onset and maintained for 4 s, followed (and preceded) by 4 s of high-frequency carriers (2000 Hz/2000 Hz). In the TIS+Cue delayed condition, the 90 Hz beat frequency preceded cue onset by 4 s and switched back to high-frequency concurrent with cue presentation, preserving identical stimulation parameters whilst temporally dissociating hippocampal TIS from the reactivation event. In the Cue only condition, no electrical stimulation was delivered. Sounds for the No-Cue condition were not administered during the nap. **c.** Representative hypnogram, illustrating the distribution of trial delivery across sleep stages for the three conditions with Cues (and therefore present during the nap). Trials were administered during NREM sleep (N2/N3). **d.** TIS electrode montage showing the four electrode positions on a representative head model. Electrode pair e1 and e2 (blue) delivered 2000 Hz, whilst electrode pair e3 and e4 (pink) switched between 2090 Hz during TIS and 2000 Hz during high-frequency carrier only. **e.** Electric field modulation magnitude metrics. **i.** TI field modulation strength (V/m) and **ii.** selectivity at the hippocampal (HP) head, body and tail, showing a significant anterior-to-posterior gradient (***p<10⁻³⁰). The hippocampal head received the strongest stimulation (0.360 ± 0.061 V/m) and higher selectivity. Values are mean ± SEM, dots represent individual participants; N = 28. **f.** Representative EEG traces during a TIS+Cue trial (top) and a Cue only trial (bottom), time-locked to cue onset (dashed line), illustrating a clear slow oscillation and sleep spindle.

TIS was delivered via two pairs of scalp electrodes positioned to target the left hippocampal head (**Fig.1d**). Both pairs delivered a 2000 Hz carrier continuously throughout stimulation windows at a 1:3 mA current ratio; shifting one pair to 2090 Hz introduced a 90 Hz frequency offset, producing a 90 Hz amplitude-modulated envelope focused at the target for a period of 4 seconds. This design ensured the two TIS conditions were matched for total stimulation exposure, including both carrier and envelope, and differed only in the envelope’s timing relative to the cue (frequency offset starts at cue onset for TIS+Cue and finishes at cue onset for TIS+Cue delayed). Subject-specific finite-element models of the TIS-induced electric field showed the expected anterior-to-posterior gradient across left hippocampal subregions^19^, consistent across field modulation magnitude metrics (TI strength, selectivity, collateral exposure; all subregions F-tests p < 10⁻³⁰, η² ≥ 0.92; (**Fig. 1e**, **Supplementary Table 1, Supplementary Fig. 1**). The hippocampal head received the strongest, most selective, and least off-target stimulation (field strength 0.360 ± 0.061 V/m, range 0.260–0.468 V/m; selectivity 1.96 ± 0.266, collateral exposure 7.91 ± 2.16%).

High-density EEG (64-channels) was recorded throughout the nap. TIS-related artefacts were effectively removed during preprocessing, with spindles and slow oscillations clearly identifiable during TIS trials (**Fig. 1f**), and power spectral analysis confirming preservation of typical 1/f sleep EEG spectra with prominent slow oscillation and spindle peaks (**Supplementary Fig. 2a**). Switching the carrier frequency to introduce or remove the 90 Hz offset produced a brief transient artifact (∼100 ms, **Supplementary Fig. 2b**) at each transition; EEG analyses for all conditions therefore excluded a conservative 250 ms on either side of every frequency transition. Cue-locked ERP morphology and oscillatory activity were preserved across conditions, confirming that TIS delivery did not contaminate the cue-evoked EEG response (**Supplementary Fig. 2c**).

TIS and TMR did not disrupt natural sleep architecture (**Supplementary Table 2**). Participants slept for 50.6 ± 13.9 minutes, with 66.8% of sleep time in N2 and 11.9% in N3. Trials were balanced across conditions and sleep stages (mean trials per condition: 40.8 ± 3.4 N2 and 11.5 ± 3.3 N3; no main effect of condition F(2,162) = 0.37, p = 0.690; or condition × sleep stage interaction: F(2,162) = 0.24, p = 0.790; **Supplementary Fig. 3, Supplementary Table 3**), ruling out sleep stage differences as a confound. These findings establish that simultaneous TIS and high-density EEG recording during sleep is feasible without compromising sleep quality while capturing brain activity during stimulation.

### Timing-dependent effects of hippocampal TIS on associative memory consolidation

We first compared performance across the three memory measures, word recognition, associative memory, and memory fidelity, before and after the nap. Performance declined over the retention interval, consistent with natural forgetting: word recognition (11.5% decline from pre- to post-nap, t(27) = −5.83, p<0.001, d = 1.10), associative memory (11.1% decline, t(27) = −7.69, p<0.001, d= 1.45), and memory fidelity (scored 0–3, with higher scores indicating better performance, showing an average decline of 1.67 to 1.27, t(26) = −3.49, p = 0.002, d = 0.67), (**Supplementary Fig. 4**). To quantify the magnitude of forgetting we computed a forgetting index defined as the difference between pre- and post-nap performance, with higher values indicating greater forgetting. Subsequent analyses examined whether this forgetting differed across stimulation conditions.

Word recognition, which can be supported by non-hippocampal processes^25,26^, showed forgetting rates that did not differ across conditions (**Fig. 2a**, binomial GLMM: χ²(3) = 4.16, p = 0.244; with forgetting rates of 22.6% ± 2.0%, 28.1% ± 2.1%, 25.3% ± 2.0%, and 24.3% ± 2.0% for TIS+Cue, TIS+Cue delayed, Cue, and No-Cue respectively).

**Figure 2.**
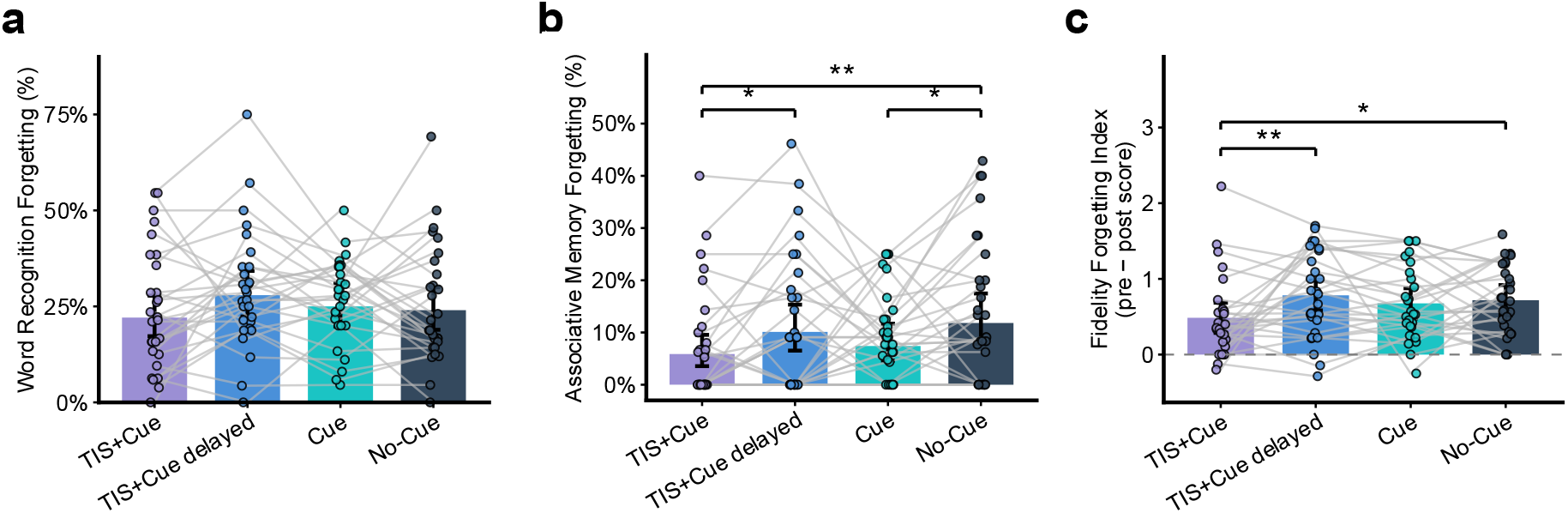
Timing effect of hippocampal TIS on memory consolidation. **a.** Word recognition forgetting (%) across the four conditions (TIS+Cue, TIS+Cue delayed, Cue, and No-Cue), where there was no significant effect of condition (χ²(3) = 4.16, p = 0.244). **b.** Associative recall forgetting (%) differed significantly across conditions (χ²(3) = 10.66, p = 0.014). Post-hoc comparisons revealed significantly less forgetting for TIS+Cue relative to both TIS+Cue delayed (*p = 0.027) and No-Cue (**p = 0.003), and significantly less forgetting for Cue relative to No-Cue (*p = 0.037). **c.** Memory fidelity forgetting index (pre − post score on 0–3 scale) differed significantly across conditions (F(3,78) = 3.03, p = 0.034). Post-hoc comparisons revealed significantly less degradation for TIS+Cue relative to both TIS+Cue delayed (**p = 0.005) and No-Cue (*p = 0.028). Values are mean ± 95% confidence intervals, dots represent individual participants; N = 28 for Word Recognition and Associative Recall and N=27 for Fidelity.

We next examined associative memory, which requires hippocampal-dependent retrieval^27^ and is sensitive to sleep-dependent consolidation^4^. Forgetting rates differed significantly across conditions (GLMM: χ²(3) = 10.66, p = 0.014; **Fig. 2b, Supplementary Table 4**), with TIS+Cue showing the least forgetting (7.7% ± 1.4%), followed by Cue (9.5% ± 1.6%), TIS+Cue delayed (12.2% ± 1.8%) and No-cue (14.7% ± 1.9%). Post-hoc comparisons revealed that TIS+Cue showed significantly less forgetting than both TIS+Cue delayed (z = −2.21, p = 0.027) and No-cue (z = −2.98, p=0.003), while the comparison with Cue alone did not reach significance (z = −0.90, p = 0.368). TIS+Cue delayed showed numerically greater forgetting than Cue alone (z = 1.33, p = 0.185), suggesting mistimed stimulation may interfere with the natural consolidation benefit of reactivation. TMR efficacy was confirmed by significantly less forgetting for cued than non-cued items (z = −2.09, p = 0.037).

We then examined whether a similar TIS timing-dependent pattern extended to memory fidelity, the richness of episodic detail. Fidelity degradation differed across conditions (LMM: F(3,78) = 3.03, p = 0.034; **Fig. 2c**, **Supplementary Table 5**), with TIS+Cue showing significantly less degradation than both TIS+Cue delayed (t = −2.86, p = 0.005) and No-cue (t = −2.24, p = 0.028), whereas TIS+Cue did not differ from Cue (t = −1.84, p = 0.070) and no other pairwise contrasts were significant (all ps > 0.30).

Finally, we asked whether electric field metrics in the hippocampal head, identical for both TIS conditions and differing only in envelope timing relative to reactivation, related to associative memory forgetting. Only selectivity showed a weak interaction with timing (F(1,26) = 4.12, p = 0.053), not present for strength or collateral (ps > 0.750). We also observed a significant main effect of field strength on forgetting (F(1,26) = 5.61, p = 0.026), absent for selectivity or collateral (ps > 0.269), **Supplementary Table 6**. Since a main effect without a timing interaction could reflect a non-specific tendency toward greater or lesser forgetting overall, we repeated the models with each participant’s Cue only forgetting rate as a covariate. The selectivity interaction was unaffected (F(1,26) = 4.12, p = 0.053; as the Cue only covariate is orthogonal to the within-participant condition contrast), with higher selectivity trending toward less forgetting in TIS+Cue (TIS+Cue: β = −18.74, SE = 10.30, t(41.7) = −1.83, p = 0.075; TIS+Cue delayed: β = 3.03, SE = 10.30, t(41.7) = 0.30, p = 0.769) **Supplementary Fig. 5**, while the strength main effect did not (ps > 0.150). TIS condition remained a significant predictor in all three models (all p ≤ 0.019), **Supplementary Table 6**.

Overall, hippocampal TIS delivered concurrently with reactivation reduced associative memory forgetting relative to both mistimed stimulation and non-cued items, a pattern mirrored in the preservation of episodic detail, demonstrating that precise temporal alignment of hippocampal stimulation is critical to preserve the consolidation benefit of memory reactivation.

### Timing-dependent enhancement of fast sleep spindles by hippocampal TIS

Having established timing-dependent behavioural effects on associative memory consolidation, we next examined whether changes in sleep electrophysiology accompanied the behavioural findings. We detected fast (12–16 Hz) and slow (10–12 Hz) spindles and SO (0.5–2 Hz) events across all EEG channels, within a 3.5-second window starting 250 ms following cue onset. Spindles (fast and slow) and SO density showed the canonical parietocentral, frontocentral and frontal topographies, respectively, which were consistent across conditions **(Supplementary Figs. 6a-c).** The same cross-condition consistency was observed for spindles and SO morphologies **(Supplementary Figs. 6d-f).**

We examined whether stimulation timing modulates the amplitude and density of spindle and SO events. Fast spindle amplitude differed significantly across conditions (channel-wise LME, cluster-corrected p<0.05) revealing two significant clusters spanning frontolateral, temporal, and centroparietal regions bilaterally (left cluster: 9 channels, cluster mass = 42.829, p<0.001; right cluster: 10 channels, cluster mass = 47.850, p<0.001; **Fig. 3a i)**. Follow-up post-hoc pairwise comparisons, limited to the channels with significant main effect of condition, revealed that TIS+Cue produced significantly greater spindle amplitude than both Cue alone (TIS+Cue: 32.5 ± 1.3 µV; Cue: 29.3 ± 1.0 µV; t(27) = −2.40, p = 0.024) and TIS+Cue delayed (29.3 ± 1.4 µV; p = 0.049), while TIS+Cue delayed did not differ from Cue (p = 0.476; **Fig. 3a ii**). Slow spindle amplitude also showed a significant main effect of condition, albeit of smaller magnitude, spanning central (4 channels, cluster mass = 16.900, p = 0.023) and right lateralised electrodes (8 channels, cluster mass = 32.365, p<0.001; **Fig. 3b i**), but post-hoc comparisons revealed no significant differences between conditions (Cue: 27.8 ± 1.1 µV; TIS+Cue: 29.2 ± 1.3 µV; TIS+Cue delayed: 29.2 ± 1.7 µV, all ps >0.35; **Fig. 3b ii**). Exploratory analysis indicated that a significant effect was only detectable within one cluster (**Supplementary Fig 7**), suggesting a weaker more regional effect than observed for fast spindles. SO amplitude differed across conditions, with a significant cluster localised over left temporal scalp regions (7 channels, cluster mass = 49.255, p<0.001; **Fig. 3c i**). However, this effect was not TIS-timing specific, with both TIS+Cue (55.8 ± 1.6 µV) and TIS+Cue delayed (57.0 ± 1.7 µV) showing greater SO amplitude than Cue alone (52.7 ± 1.6 µV; Cue vs TIS+Cue: t(27) = −2.50, p = 0.019; Cue vs TIS+Cue delayed: t(27) = −2.40, p = 0.024), while the stimulation conditions did not differ from each other (t(27) = −1.36, p = 0.186; **Fig. 3c ii**).

**Figure 3.**
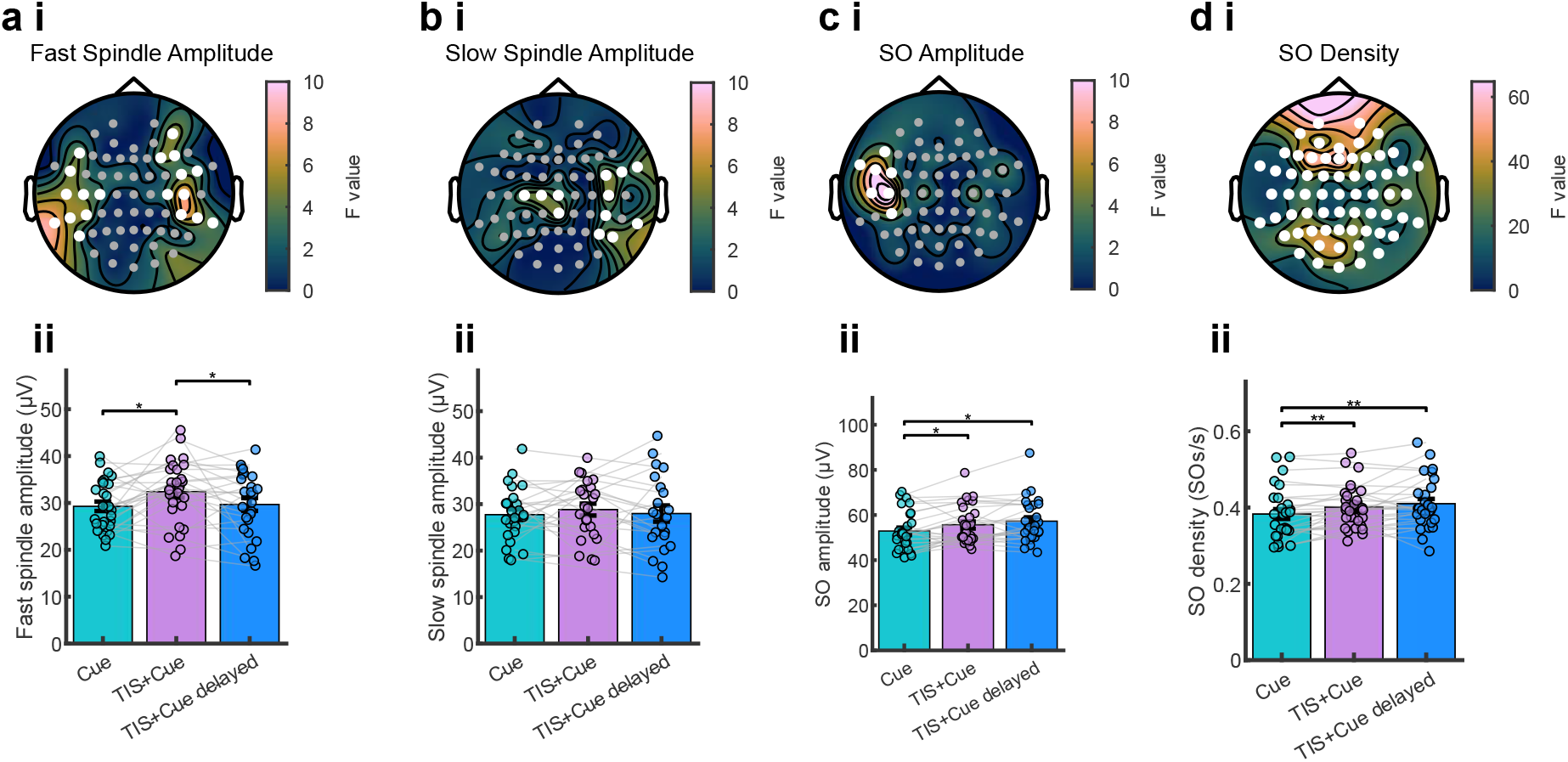
TIS-timing modulates spindle and slow oscillation (SO) dynamics. For each metric derived from the corresponding electrophysiological event (fast spindle: 12-16 Hz, slow spindle: 10-12 Hz, and SOs: 0.5-2 Hz), the top row shows topographical distribution of F-statistics from channel-wise LMEs (colourbar) computed from data from a 3.5 second window starting 250 ms after cue onset, covering the memory reactivation period. Each channel is represented by a grey dot. White dots indicate significant channels belonging to a cluster surviving cluster-based permutation correction (p<0.05). Bottom plots display post-hoc pairwise comparisons within all channels from the significant clusters. **a i.** Topoplot of fast spindle amplitude across conditions (Cue, TIS+Cue, TIS+Cue delayed) showing significant clusters on the right and left hemispheres spanning frontolateral, temporal, and centroparietal regions, and **ii** showing that TIS+Cue had significantly greater fast spindle amplitude than both Cue (*p = 0.024) and TIS+Cue delayed (*p = 0.049). **b i.** Slow spindle amplitude with two clusters showing a significant effect of condition in central and right-lateralised electrodes, **ii.** but no significant post-hoc pairwise comparisons within the significant cluster (all p > 0.35). **c i.** SO amplitude displaying one signficant cluster in left-temporal channels, and **ii.** Showing that both TIS+Cue (*p = 0.019) and TIS+Cue delayed (*p = 0.024) had greater SO amplitude than Cue alone. **d i.** SO density (Gamma GLME) revealed a cluster spanning all head electrodes, and **ii** shows that both TIS+Cue (**p = 0.004) and TIS+Cue delayed (**p = 0.010) have greater SO density than Cue alone. Values are mean ± SEM, dots represent individual participants; N = 28 throughout.

Neither fast nor slow spindle density differed significantly across conditions (channel-wise Gamma GLME, cluster-corrected p<0.05; **Supplementary Fig. 8**), indicating that TIS modulated the amplitude but not the rate of spindle events. SO density differed significantly across conditions in the whole-scalp (**Fig. 3d i**). Post-hoc comparisons showed that both TIS conditions produced significantly higher SO density than Cue alone (Cue: 0.383 ± 0.013 SOs/s; TIS+Cue: 0.401 ± 0.011 SOs/s; t(27) = −3.12, p = 0.004; TIS+Cue delayed: 0.410 ± 0.013 SOs/s; t(27) = −2.77, p = 0.010), while TIS+Cue and TIS+Cue delayed did not differ from each other (t(27) = −1.21, p = 0.238; **Fig. 3d ii**).

This pattern indicates that hippocampal stimulation broadly enhances SO activity during reactivation, while selective amplification of fast spindles requires precise temporal coordination between stimulation and memory reactivation.

### Mistimed stimulation disrupts the relation between coupling rate and consolidation

We next examined the coordination between SOs and fast spindles as these have been associated with temporal coordination during memory reactivation^11,12^. We focused on two metrics: SO-spindle coupling rate, defined as the percentage of spindle events temporally co-occurring with a SO, and coupling strength, indexed by the mean vector length (MVL) of spindle peak phases relative to the SO cycle.

Coupling strength was highest over frontocentral, central, and centroparietal channels in all conditions with no significant condition effect on MVL (ps > 0.3; **Fig. 4a**); this topography was used to define a region of interest for subsequent phase analyses (see Methods). Rayleigh tests confirmed significant non-uniform phase distributions in all three conditions (TIS+Cue: MVL = 0.777, z = 16.91; TIS+Cue delayed: MVL = 0.692, z = 13.39; Cue: MVL = 0.637, z = 11.35; all ps <0.001), with spindles preferentially occurring on the rising phase of the SO approaching the upstate peak (TIS+Cue: −37.9°; TIS+Cue delayed: −52.3°; Cue: −34.5°; 0° = SO upstate peak; **Fig. 4b).** The preferred coupling phase did not differ significantly between conditions (Watson-Williams test: F(2,81) = 0.842, p= 0.434), indicating that TIS did not shift the timing of spindle occurrence relative to the SO cycle. Similarly, for SO-spindle coupling rate there was no significant effect of condition (ps > 0.2, **Fig. 4c**).

**Figure 4.**
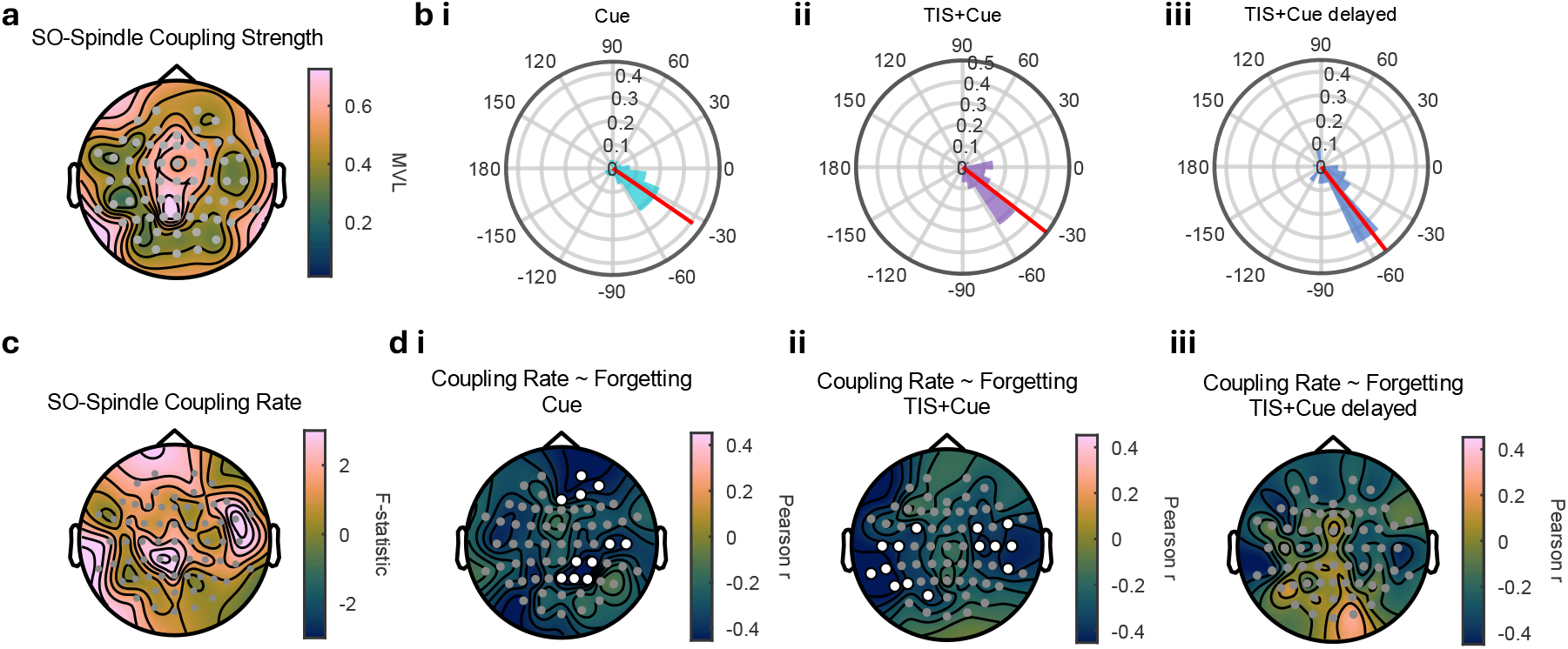
Effect of TIS timing on SO-spindle coupling rate and strength. **a.** Topographical distribution of SO-spindle coupling strength (mean vector length, MVL), averaged across the three conditions. Coupling strength was maximal over frontocentral, central, and centroparietal regions; no significant main effect of condition (channel-wise LME, cluster-based permutation correction p<0.05). **bi–iii.** Rose plots illustrating the preferred SO phase of spindle coupling for Cue (i), TIS+Cue (ii), and TIS+Cue delayed (iii). Individual participant mean phase vectors (coloured wedges) and the group mean resultant vector (red line) are shown. Rayleigh tests confirmed significant non-uniform phase distributions in all three conditions (all ps < 0.001), with spindles preferentially occurring on the rising phase of the slow oscillation approaching the upstate peak. The preferred coupling phase did not differ significantly across conditions (Watson-Williams test: p = 0.434). **c.** Topographical distribution of F-statistics from channel-wise LMEs comparing SO-spindle coupling rate across conditions. No clusters survived cluster-based permutation correction. **di–iii.** Topographical distribution of Pearson r values from channel-wise correlations between SO-spindle coupling rate and associative forgetting for Cue (i), TIS+Cue (ii), and TIS+Cue delayed (iii). Significant negative associations between coupling rate and forgetting were observed in the Cue and TIS+Cue conditions; no clusters survived correction in TIS+Cue delayed. White dots indicate electrodes belonging to clusters surviving cluster-based permutation correction (p < 0.05). N = 28 throughout.

Although we did not find a significant effect of condition in these metrics, SO–spindle coupling has been linked to consolidation; we therefore examined whether it related to individual differences in associative memory forgetting. We computed per condition channel-wise Pearson correlations between coupling strength (MVL) and coupling rate and associative memory forgetting across all EEG channels. While for MVL there were no significant clusters surviving permutation correction in any condition **(Supplementary Fig. 9)**, coupling rate showed significant associations with forgetting. In the Cue condition, two clusters, over right centroparietal (7 channels, cluster mass = 17.31, p = 0.027) and right prefrontal scalp (4 channels, cluster mass = 9.44, p = 0.050), showed that higher coupling rate during memory reactivation was associated with less forgetting (**Fig. 4d i, Supplementary Fig. 10i)**. Similarly, significant correlations were present in the TIS+Cue condition (two significant clusters, both reflecting negative associations between coupling rate and forgetting: one over left temporocentral-parietal scalp, 8 channels, cluster mass = 19.87, p = 0.025, and one over right frontocentral-temporal scalp 6 channels, cluster mass = 13.43, p = 0.039) (**Fig. 4d ii, Supplementary Fig. 10ii).** No such associations were observed in the TIS+Cue delayed condition (**Fig. 4d iii**). The coupling-forgetting relationship was thus present when reactivation occurred with or without coincident stimulation, but absent when stimulation was mistimed, consistent with a timing-dependent pattern as seen for behaviour and fast spindles.

### Distributed EEG dynamics link hippocampal TIS to associative memory

Since TIS’s effects on brain activity and their relationship to behaviour are likely distributed across multiple measures beyond the discrete oscillations and their coupling investigated in the previous sections, we tested whether a wider ensemble of continuous EEG features mediated the effect of TIS on associative memory accuracy (mediation model, **Fig. 5a**). To this end, we extracted 51 features, quantified regardless of whether a discrete oscillation was detected, allowing trial-level analysis across the full set. These comprised four spectral measures (spindle-band power, spindle peak frequency, theta power, spectral slope), one envelope measure (sigma burst amplitude), and three phase–amplitude coupling (PAC) measures (SO–spindle, delta–spindle, and their difference), each averaged within six cortical regions (frontal, central, left/right temporal, posterior, occipital), plus three frontal-to-central (F→C) PAC features linking frontal SO or delta phase to central spindle amplitude. All features reflected post-minus pre-cue change, focusing the analysis on the memory reactivation period, and were z-scored to a common scale. To mitigate collinearity, hierarchical clustering identified 16 maximally representative, minimally correlated features (cluster threshold |r| ≤ 0.5; **Supplementary Fig. 11**), which were carried forward to the mediation analysis.

**Figure 5.**
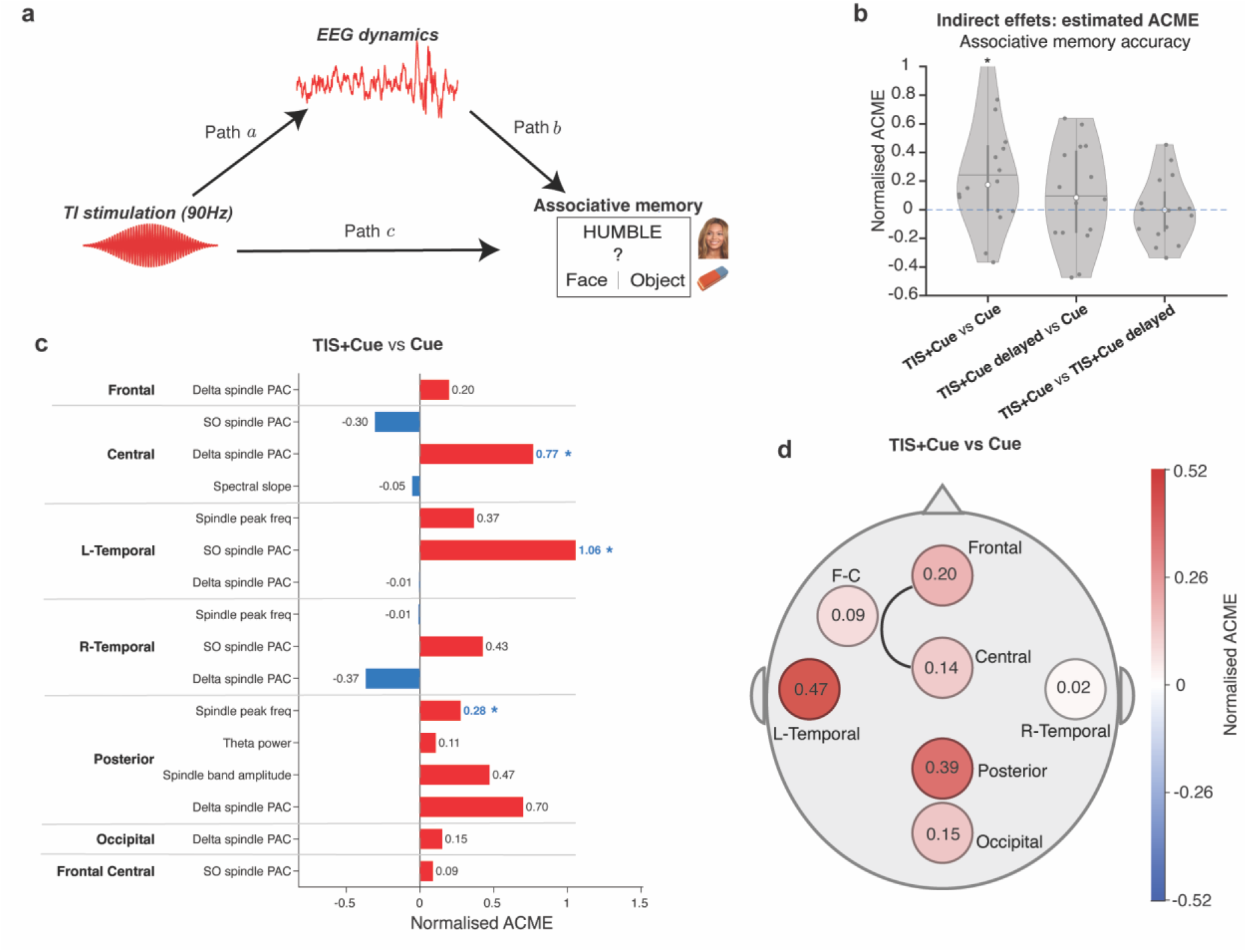
TIS delivered with Cue modulated the associative memory performance via a set of EEG features. **a.** Mediation framework: TIS can impact associative memory by modulating EEG dynamics (paths a and b, the indirect effect modelled) or through other mechanisms (path c). **b.** Distribution of per-feature normalised average causal mediation effect (ACME) values across the 16 features retained for modelling. The dots show each feature’s value; violins show the kernel density (shape), median (non-filled circle), and upper and lower 25% quantiles (grey bar). Dashed line at zero ACME. ‘*’ indicates p<0.05 for two-sided one-sample t-test. **c.** Per-feature CoV grouped by region (Frontal, Central, left L-Temporal, right R-Temporal, Posterior, Occipital, and cross-channel Frontal-to-Central, F-C). Colours indicate positive (red) and negative (blue) values. ‘*’ indicates p<0.05 for Surrogate testing. **d.** Topological map of the averaged ACME values per region. Colour indicates the average ACME values for the features within that region.

We first examined how TIS modulated these features (path-a effects; **Supplementary Table 7, Supplementary Fig. 12**). The central aperiodic slope increased (became steeper) under TIS delivered with Cue relative to Cue alone (TIS+Cue: β = 0.12, t(3176) = 3.47, pFDR = 0.008) and TIS before Cue (TIS+Cue delayed: β = 0.202, t(3073) = 5.86, pFDR = 8.4 × 10⁻⁸).

Posterior spindle peak frequency decreased under TIS+Cue relative to Cue alone (β = −0.096, t(3173) = −2.86, pFDR = 0.034). See **Supplementary Table 7** for full statistics.

We estimated the average causal mediation effect (ACME) of TIS+Cue on associative memory accuracy through each feature using a joint mediator model comprising all 16 features, such that each feature’s ACME reflects its independent contribution conditioned on all remaining features. We found that TIS+Cue exerted a positive indirect effect on associative memory through 11 of 16 features compared to Cue alone (normalised ACME values across features mean ± SD = 0.24 ± 0.38, median = 0.17; two-sided one-sample t-test against 0, p = 0.02; **Fig. 5b**). This indicates a coordinated indirect effect distributed across the feature ensemble rather than carried by any single feature. This pattern was absent for TIS+Cue delayed versus Cue only and for TIS+Cue versus TIS+Cue delayed (mean normalised ACME ± SD = 0.10 ± 0.34, median = 0.09 and −0.001 ± 0.22, median = −0.001; two-sided one-sample t-test: p = 0.28 and p = 0.99, respectively). Although no feature reached significance under the model’s own quasi-Bayesian ACME p-values, comparing each feature’s ACME against a within-subject surrogate null (1000 permutations, one-sided), three features showed nominal evidence of positive unique mediation: posterior spindle peak frequency (p = 0.013), left temporal SO–spindle coupling (p = 0.030), and central delta–spindle coupling (p = 0.044; **Fig. 5c, Supplementary Table 8**). Examining the mediator–outcome associations within the joint model, only posterior sigma burst amplitude was reliably associated with associative memory accuracy after correction (β = 0.13, pFDR = 0.038); this feature was, however, weakly modulated by TIS **(Supplementary Table 8)** and thus contributed little to the mediated effect. We further inspected the ACME per brain region, revealing a left-lateralised pattern (**Fig. 5d).** The left temporal region showed the highest mean normalised ACME (0.47), driven predominantly by SO–spindle PAC, followed by posterior (0.39; delta–spindle coupling and sigma burst amplitude) and frontal regions (0.20).

In sum, mediation analyses revealed a coordinated, left-dominated indirect pathway through which TIS+Cue modulates associative memory accuracy, although the modest magnitude of this mediated component suggests the presence of additional mechanistic pathways not identified by the scalp EEG.

## DISCUSSION

Here, we showed that non-invasive hippocampal stimulation via TIS, delivered concurrently with targeted memory reactivation during NREM sleep, reduces associative memory forgetting relative to both mistimed stimulation and non-cued items, with a parallel preservation of episodic detail. This timing-dependent behavioural benefit was accompanied by timing-specific enhancement of fast spindle amplitude, a timing-dependent relationship between SO-spindle coupling rate and memory and a left-lateralised ensemble of EEG mediators linking stimulation to memory. Together, these findings constitute the first demonstration that non-invasive deep brain stimulation can modulate sleep-dependent memory consolidation in humans, with efficacy contingent on precise temporal alignment between hippocampal stimulation and memory reactivation.

The hippocampus is thought to play a central role in binding and retrieving associative memories^27^. Accordingly, the behavioural effects were specific to memory that required retrieval of the encoded association, showing less forgetting under TIS+Cue than under mistimed stimulation or non-cued items; a pattern also present in memory fidelity. Although fidelity draws on neocortical, semantic, and perceptual representations beyond the hippocampus^28^, its convergence with associative memory points to stimulation acting on a shared process. No such effect was observed for word recognition. Although item recognition can draw on hippocampal recollection, it can be supported by processes not requiring explicit associative retrieval^25,26^. This specificity is consistent with stimulation engaging hippocampal mechanisms supporting consolidation during sleep rather than producing non-specific arousal or sensory effects. This also aligns with the effects of stimulation acting on the memory reactivation processes induced by TMR, which primarily benefits recall and associative memory over recognition^21^. Our findings contrast with a recent study reporting improved recognition but not associative memory when intracranial stimulation was applied to the prefrontal cortex during sleep^17^. That prefrontal stimulation benefited item recognition whereas hippocampal stimulation affected associative memory and fidelity points to anatomical specificity of the effect rather than a generic consequence of precisely timed stimulation during sleep^27^. Yet, the timing of the stimulation is critical. TIS+Cue delayed produced greater forgetting than TIS+Cue despite being matched for total carrier and envelope exposure, electrode configuration, and current amplitude, differing only in whether the 90 Hz envelope coincided with the auditory reactivation cue. This finding parallels the same intracranial deep brain stimulation study^17^, where stimulation delivered outside the SO active window, detected in the medial temporal lobe failed to improve, and sometimes degraded memory, and extends the principle that timing relative to reactivation determines consolidation outcome from invasive to non-invasive electrical stimulation. Associative memory forgetting under TIS+Cue was numerically, but not significantly, lower than under Cue alone. Comparable patterns have been reported in within-subject phase-locked TMR studies, where SO-upstate outperformed SO-downstate cueing but shows no consistent additional benefit over uncued items^29,30^, possibly reflecting a ceiling on the benefit reactivation-timing manipulations can add in within-subject designs restricted to one sleep episode. It is possible that this effect would require more extensive stimulation, e.g., a full night with more reactivation opportunities to emerge, or the delivery of only one condition per sleep episode.

What is the possible mechanism by which concurrent hippocampal TIS and memory reactivation selectively enhanced fast spindle amplitude? The 90 Hz TIS envelope falls within the human hippocampal ripple band^5–7^, raising the possibility that subthreshold stimulation at this frequency engages or amplifies ongoing ripple activity during reactivation, with upstream thalamocortical consequences manifesting as increased fast spindle amplitude. Fast spindles are preferentially coupled to hippocampal sharp-wave ripples to support hippocampal– neocortical memory transfer^6,13,16,31^. That TIS enhanced fast but not slow spindles concords with their preferential role in hippocampus-dependent consolidation^12^. We therefore tentatively interpret the amplitude effect as a scalp-level expression of enhanced fast-spindle-ripple coordination during reactivation, an account supported by its specificity to TIS+Cue, the condition that produced the better memory outcomes. However, whether TIS directly modulates hippocampal ripples cannot be resolved from scalp EEG.

In contrast, SO amplitude and SO density were increased by both TIS conditions relative to Cue alone, with the amplitude effect localised to the left temporal region, consistent with our left hippocampal target. This SO effect is unlikely related to behaviour, as it was comparable across TIS conditions despite their differing forgetting rates, consistent with evidence that SO amplitude and density are not reliable correlates of consolidation^12^, and with a wider literature in which non-invasive stimulation reliably enhances SOs without consistent memory benefit^32^. What can then explain the timing-independent modulation of SOs by TIS? Two non-exclusive accounts remain compatible with our data: carry-over effects from the 90 Hz stimulation in the TIS+Cue delayed condition, or SOs simply being more readily modulated. As large-amplitude, globally distributed cortical events, SOs present a broader target for non-specific diffuse perturbation by the kHz carrier itself, independent of reactivation-locked engagement. However, a recent TIS study found that 1 Hz-modulated TIS enhanced SO activity whereas the carrier alone did not^33^ Yet their carrier was substantially higher than ours (15 vs 2 kHz) and whether carrier frequencies differ in effects remains to be established^34^.

What accounts for the timing-dependent modulation of the SO-spindle coupling-memory relationship? SO-spindle coupling rate was associated with individual differences in associative memory forgetting in TIS+Cue and Cue but not in TIS+Cue delayed, indicating that this coupling-memory relationship, present with cued or coincident stimulation, was disrupted when stimulation was mistimed. This is consistent with evidence linking spindle-SO co-occurrence to declarative memory consolidation^6,35^ and with SO–spindle coupling providing a temporal gate for hippocampal-neocortical transfer^14,35^. By the same principle, engaging hippocampal circuits outside this window may interfere with this gate. The topography of this coupling however differed between conditions. In Cue alone the association was frontocentral and centroparietal, consistent with more commonly reported frontal-to-frontocentral localisation^12,35^ but see^23^. In TIS+Cue, left temporocentral-parietal involvement emerged alongside this pattern. It is possible that by targeting the left anterior hippocampus TIS preferentially engaged left-lateralised circuits. Potential support for this comes from evidence that unilateral TMR produces hemisphere-specific effects on SO-spindle coupling^36^. Our design cannot however establish whether this topography would differ had we targeted the right hippocampus. Across all conditions, spindles preferentially occurred on the rising phase of the slow oscillation approaching the up-state peak, with no significant difference between conditions, in agreement with the established phase relationship between these events during NREM sleep^6,35^. This indicates that the timing-dependent effects of TIS operated on the rate and behavioural relevance of coupling rather than on its preferred phase.

The mediation analysis allowed testing whether, and through which EEG features, the effect of TIS on memory was expressed. We found that this was conveyed through a coordinated ensemble of features in which spindle and SO-spindle coupling measures were the most prominent mediators, with SO-spindle coupling the strongest. This agrees with findings that sleep-dependent reactivation is distributed across many features rather than carried by any one correlate^37^. That SO-spindle coupling emerged as a leading mediator here but not in the discrete event-based analysis may reflect the greater sensitivity of continuous time-frequency measures of sigma activity relative to thresholded spindle detection^38^. Notably, the mediated effect was of higher magnitude in the left temporal region, converging with the additional left hemispheric coupling-forgetting topography, and the left-hemisphere target. That this mediated pathway nonetheless accounted for only part of the total behavioural effect might be expected, since scalp EEG cannot resolve the hippocampal sharp-wave-ripple dynamics most directly implicated in associative consolidation.

While we observed timing-dependent effects on behaviour, we did not observe a clear relationship between field metrics and associative memory performance. Only selectivity showed a weak association with the timing manipulation, consistent with a possible contribution of field focality. Field modulation strength related to forgetting independently of timing, but this association did not survive control for individual differences in overall forgetting. In our previous study we found a weak relationship between BOLD signal in the hippocampus and field modulation strength^19^. In a tACS study examining both properties, field spatial precision independently predicted the physiological aftereffect, whereas field strength alone showed only a weak, spatially inconsistent correlation^39^. Measures reflecting activity at the target area may simply be more sensitive to field metrics than distal behavioural outcomes, but resolving this will require a direct hippocampal readout, which the present study does not provide.

The current design does not permit disambiguation of whether TIS acted via hippocampal ripple augmentation, spindle-mediated thalamocortical effects, or a combination of both. Concurrent intracranial recordings will be necessary to resolve this question and to establish whether selectivity and field modulation strength produce proportionally stronger effects on local activity. Our sample comprised healthy young adults, and whether the approach generalises to populations with impaired sleep-dependent consolidation remains to be examined. Notably, these effects were achieved with brief, 4 s bursts of TIS delivered per reactivation even, roughly eightfold shorter than previous fMRI block designs and around two orders of magnitude shorter than average session duration reported across published human TIS studies^40^. TIS was nonetheless delivered in a fixed temporal relationship to the auditory cue. Future studies delivering TIS in a closed-loop fashion tuned to natural windows of reactivation may yield stronger consolidation benefits.

Sleep-dependent consolidation is disrupted across many brain disorders, including Alzheimer’s disease, schizophrenia, and depression. The present findings demonstrate that TIS can non-invasively modulate hippocampal contributions to sleep-dependent consolidation in healthy humans, with effects contingent on temporal coordination with reactivation. These findings open tractable avenues for non-invasive hippocampal interventions in populations where consolidation is compromised.

## METHODS

### Participants

We recruited 42 healthy adults through posters, emails, and the School of Psychology participant recruitment system. Two participants were excluded after the MRI session due to excessive head motion, and one participant withdrew from the study after the MRI session. Following the nap session, we excluded 11 additional participants: nine could not achieve N2 nor N3 sleep, one failed to attend the scheduled session and one because of technical difficulties with the equipment. Our final sample was comprised of 28 right-handed participants (15 males, 13 females) with mean age of 24.8 years (SD = 6.0 years, range 18-37 years). All participants had completed or were pursuing higher education qualifications. We required participants to be aged 18-40 years, with no self-reported history of sleep, neurological or psychiatric illness, not taking medications affecting sleep or cognitive function, not pregnant, not exceeding alcohol intake of 28 units per week, and be free from contraindications for MRI or transcranial electrical stimulation. The sample size was determined based on reported effect sizes from previous studies showing effects of non-invasive brain stimulation on memory performance (0.25 – 0.7)^19,41^. To detect the smallest effect size (0.25) using a repeated measures ANOVA (1 group × 4 measurements) with an α-level of 0.05 and power of 0.8, the estimated sample size was 24 participants (G*Power), which is consistent with sample sizes used in previous human sleep and memory studies^23,42–44^. We recruited a larger sample based on conservative estimates of attrition due to possible insufficient sleep or technical issues.

The study protocol was approved by the University of Surrey Ethics Committee (FHMS 22-23 123 EGA / ERM 873). All participants provided written informed consent, and procedures were in accordance with the Declaration of Helsinki. Participants received financial compensation or course credit via the School of Psychology research participation scheme, for their time.

### Experimental Procedure

The study included two sessions on separate days: an MRI session (at the Combined Universities Brain Imaging Centre (CUBIC), see MRI Data Acquisition) followed by the sleep session (at the Surrey Sleep Research Centre). Participants were instructed to maintain regular sleep schedules at least 3 days prior to the sleep session, avoid alcohol for 24 hours, and abstain from caffeine on the day.

The sleep session occurred in the afternoon, starting at ∼1 pm, with the following order of events: 1) TIS setup and test of participants’ sensations and comfort (see Temporal Interference Stimulation); 2) EEG setup; 3) Registration of TIS electrode positions; 4) Memory task (image-pair associates) composed of familiarisation and encoding phases followed by an immediate memory recall test (see Memory Task); 5) Assessment of subjective sleepiness using the Karolinska Sleepiness Scale (KSS); 6) Lights off and 90-minutes sleep opportunity; 7) Lights on and KSS to assess post-sleep alertness; 8) Questionnaires for subjective experience assessing stimulation-related sensations, sleep quality, dream experience, and awareness of auditory cues during sleep; 9) Post-nap memory test (conducted at least 20 minutes after waking to minimise sleep inertia effects).

Procedures in the sleep session were performed in a temperature-controlled, sound-attenuated, windowless sleep laboratory. Researchers monitored participants throughout the sleep opportunity through real-time monitoring of EEG activity. We terminated the sleep opportunity after 90 minutes or if the participant spontaneously woke up and could not return to sleep within 20 minutes. Stimulation started whenever participants reached stable N2 or N3 sleep (lasting at least four consecutive 30-second epochs). If participants showed signs of arousal, stage transitions to N1 or REM, or excessive movement artefacts, stimulation was stopped and restarted after stable N2 or N3 sleep returned. The order of stimulation was counterbalanced across participants.

### Memory Task

A paired associative memory task was chosen because these types of tasks are sensitive to the effects of sleep and have been commonly used in targeted memory reactivation studies^23,45–47^. Each pair was composed of a word and an image of an object or a famous face. Objects and faces were chosen because we have previously shown that TI stimulation can modulate memory performance in a face-name task, and both faces and objects engage medial temporal lobe structures including the anterior hippocampus.

Word stimuli comprised 220 adjectives selected from the MRC Psycholinguistics Database (http://websites.psychology.uwa.edu.au/school/MRCDatabase/uwa_mrc.htm) filtered by number of letters (3–8), Kucera-Francis written frequency (minimum 20), and syllables (1–4). Of these, 120 adjectives were used for the main task, 80 served as novel foils for retrieval testing (40 pre-nap, 40 post-nap), and 20 were reserved as control cues during the nap. All words were converted to audio files using Google Text-to-Speech (British English female voice, 48 kHz sampling rate). Image stimuli included 60 face images (famous people from open-source databases) and 60 object images from the MIT Unique Objects database (http://olivalab.mit.edu/MM/uniqueObjects.html) depicting common everyday items. All images were presented on a plain grey background. The task was composed of a familiarisation phase, encoding and two retrieval phases (pre-nap and post-nap). Before the encoding participants completed a short practice session composed of three trials to ensure they understood the task instructions and response requirements. The task was designed using PsychoPy (version 2023.2.3) and presented on a 27-inch monitor (2560 × 1440 resolution, 119 Hz), with auditory cues delivered binaurally through insert earphones (SE215, Shure, United States). Response selections were done with a computer mouse.

### Familiarisation

The familiarisation phase at the beginning of the experiment was included to facilitate recognition of the images and learning of the adjective-image pairs in the main encoding phase. Participants viewed all 120 images in randomised order. Each image appeared for 4 s followed by a 1.5 s fixation cross, and participants categorised each image as “face” or “object” using mouse click.

### Encoding

Participants were presented 120 word-image associations in four blocks of 30 pairs each (15 adjective-face, 15 adjective-object). Each trial began with a 1.5 s fixation cross, followed by simultaneous presentation of the adjective (displayed as text above the centrally-positioned image) and its corresponding spoken word for 4.5 s. Participants were instructed to form a vivid mental image or story linking the word and image.

### Retrieval / Memory test

Memory retrieval was assessed pre- and post-nap using identical procedures. Each test included the 120 words presented during encoding intermixed with 40 novel foil words, divided into four blocks. Different foil sets were used pre- and post-nap to prevent foil memorisation. Each trial started with a fixation cross (1.5 s ± 0.1 s), followed by presentation of the word (text and audio). Participants first indicated whether the word was “old” (i.e., present at encoding) or “new” (foil, i.e., not present at encoding). For “new” responses, the trial ended immediately. For “old” responses, participants then indicated whether the word was paired with a face or object, provided a confidence rating (1-5 scale; 1 being unsure and 5 being very confident), and gave a verbal description of the associated image which was recorded using a microphone. For trials where the category was correctly recalled, participants were able to correctly describe the image on the majority of occasions (mean ± SD: 78.1 ± 20.9%), demonstrating that the category responses reflected veridical memory.

### Targeted Memory Reactivation (TMR)

Adjective-image pairs correctly recalled during the pre-nap test (i.e., when the adjective was correctly recognised as “old” and the associated image category correctly recalled) were equally distributed across four conditions: TIS+Cue, TIS+Cue delayed, Cue only, and No-Cue. Items were allocated proportionally by category (faces and objects). For example, if a participant recalled 60/120 pairs, 15 were assigned to each condition, with faces and objects distributed equally across conditions and any remainders randomly assigned. This ensured that the baseline category recall performance was balanced between conditions. Participants recalled on average 65/120 pairs (SD = 14.3, min = 40, max = 102), yielding approximately 10–26 items per condition per participant (mean = 16).

During the nap, three conditions involved replaying the associated adjective sounds: Cue only (in which the sound cue was presented without stimulation), TIS+Cue and TIS+Cue delayed conditions (see Stimulation Parameters for description of the TIS conditions). Non-Cued items were not presented during the nap. Additionally, foil sounds (adjectives not associated with learned items) were interspersed without stimulation at 50% of the cued sound frequency. The three cued conditions were presented in blocks in a pseudo-randomized order with the objective of distributing conditions evenly across the nap, avoiding more than three consecutive blocks of the same condition, and ensuring balanced representation during N2 and N3 sleep stages. Blocks cycled repeatedly while participants maintained stable N2 or N3 sleep. There were no differences in the distribution of trials between TIS+Cue, TIS+Cue delayed and Cue only across sleep stages (main effect of condition: F(2,162) = 0.37, p = 0.69; condition × sleep stage interaction: F(2,162) = 0.24, p = 0.79; **Supplementary Fig. 3)**

### Temporal Interference Stimulation (TIS)

TIS was delivered using an eight-channel device generating independent fully differential sinusoidal currents (TIBS-R V3.0, TI Solutions AG, Zurich, Switzerland), with synchronised, galvanically isolated channels, constant current output, automatic impedance monitoring, and real-time waveform recording. Two pairs of stimulation electrodes (self-adhesive TENS 1.5 cm diameter) were positioned as in Violante et al. (2023)^19^: two over the left temporal region (e1 anterior, e3 posterior, near the ear) and two over the right hemisphere (e2 anterior, e4 posterior), with e1–e2 forming one pair and e3–e4 the second pair. Positioning formulae based on head size are provided in Supplementary Materials. Prior to application, scalp sites were cleaned with NuPrep exfoliating gel (Weaver and Company, USA), followed by alcohol wipes and gauze. Electrodes were affixed using Ten20 conductive paste (Weaver and Company, USA) and secured with microporous surgical tape. Initial impedances were below 5 kΩ, measured using a portable impedance checker (EL-CHECK, BIOPAC Systems, Inc., USA). Throughout the recordings impedance values were kept low (mean = 1.64, SD = 0.43).

### Stimulation Parameters

Current amplitudes were set at a 1:3 mA ratio (1 mA for e1-e2, 3 mA for e3-e4, peak-to-baseline) to target the left anterior hippocampus / hippocampal head^19^. Stimulation was organised into blocks with 5 s ramps applied only at block boundaries to minimise EEG artefacts and potential sensory perception associated with current changes. Within blocks, high-frequency currents (2000 Hz/2000 Hz) were maintained continuously, switching to the target 90 Hz beat frequency (2000 Hz/2090 Hz) according to the timing condition. This approach served multiple purposes: (1) reducing EEG artefacts from repeated ramp transitions that could interfere with sleep stage detection and analysis, (2) minimising possible cutaneous sensations associated with current onset/offset that might disrupt sleep, and (3) maintaining consistent electrical stimulation across TIS conditions to isolate timing-specific effects. In the TIS+Cue condition, each 4 s trial delivered 90 Hz beat frequency starting simultaneously with sound cue onset, followed immediately by 4 s of high-frequency baseline (2000 Hz/2000 Hz at identical amplitude). In the TIS+Cue delayed condition, 90 Hz beat frequency preceded the sound cue by 4 s, then switched back to high-frequency baseline concurrent with cue presentation. This design enabled precise temporal control over TIS delivery relative to cue presentation, allowing us to investigate whether simultaneous versus sequential TIS+Cue timing differentially modulates memory reactivation and consolidation, whilst the Cue only condition (auditory cue without electrical stimulation) controlled for non-specific effects of the cue presentation itself.

Stimulation parameters were pre-programmed using the device’s Scripter software (Python interface). During sleep, PsychoPy (version 2023.2.3, Python 3.7) controlled stimulus timing by sending TTL triggers via a National Instruments USB-6343 DAQ device to a trigger converter box (TRIG-EOC, TI Solutions AG), which initiated the pre-programmed stimulation protocols. The stimulator was located in the control room, with stimulation cables routed into the sleep room through a sound-attenuated porthole window.

### Stimulation Titration, Sensation and Side Effects

Before each experiment, we individually titrated stimulation intensity to establish perceptual thresholds. We delivered conventional transcranial alternating current stimulation separately to each electrode pair, starting at 0.1 mA and increasing in 0.1 mA steps, followed by high-frequency temporal interference (2000 Hz/2090 Hz) starting at 1 mA and increasing in 0.5 mA steps until participants reported sensations or reached 3 mA maximum. At each step, participants verbally reported sensation presence and quality (tingling, itching, warmth, pressure), which were recorded in Microsoft Excel. After the experiment, participants completed a questionnaire rating the intensity and duration of possible side effects from 0 (none) to 5 (severe), including pain, burning, warmth/heat, itchiness, pinching, metallic taste, fatigue, trouble concentrating, sleepiness, headache, mood change, and any other perceived effects. Detailed descriptions of perceptual thresholds and side effects are provided in **Supplementary Tables 9 and 10**s.

### MRI

Data were acquired on a 3T Siemens Trio Tim scanner using a 32-channel head coil (Siemens, Erlangen, Germany). T1-weighted anatomical images were acquired using a magnetisation-prepared rapid gradient echo (MPRAGE) sequence with 0.8 mm isotropic voxels (TR = 2300 ms, TE = 3.2 ms, TI = 1100 ms, flip angle = 11°, field of view 256 × 256 mm, matrix = 320 × 320, 224 slices, GRAPPA acceleration factor = 2). FLAIR images were also acquired using a 3D T2-weighted SPACE sequence with 0.8 mm isotropic voxels (TR = 4000 ms, TE = 392 ms, TI = 1800 ms, flip angle = 120°, field of view 256 × 256 mm, matrix = 320 × 320, 224 slices, GRAPPA acceleration factor = 2). Diffusion-weighted images (DWI) were acquired using a spin-echo echo-planar imaging sequence, 2 mm isotropic voxel, TR = 8300 ms, TE = 90 ms, flip angle = 90°, field of view 216 × 216 mm², matrix = 108 × 108, 64 slices, GRAPPA acceleration factor = 2, partial Fourier = 6/8. A full diffusion-weighted acquisition was obtained in the anterior-posterior phase-encoding direction comprising 64 diffusion-encoding directions at b = 1000 s/mm² plus one non-diffusion-weighted (b = 0 s/mm²)) volume. An additional b = 0 s/mm² volume was acquired in the posterior-anterior phase-encoding direction. These matched phase-encoding pairs were used for susceptibility-induced geometric distortion correction.

### Electric Field Modelling

To characterise the individualised hippocampal electric field delivered by TIS subject-specific finite element head models were constructed and simulated. DWI data were preprocessed using MRtrix3 (v3.0.4). Raw data were converted to .mif format preserving gradient information. Thermal noise was removed using dwidenoise and Gibbs ringing artefacts were corrected using mrdegibbs. Motion, eddy current, and susceptibility-induced distortions were corrected using dwifslpreproc, which internally calls FSL’s topup and eddy using the anterior-posterior and posterior-anterior b = 0 s/mm² pair for distortion correction (total readout time = 0.039 s). Cortical reconstruction and subcortical segmentation were performed using FreeSurfer (v7.4.1) recon-all, with the FLAIR image supplied using the - T2pial flag. Hippocampal subfield segmentation was performed using FreeSurfer’s hippocampal subfield module (segmentHA_T1.sh), producing separate volumetric labels for the left hippocampal head, body, and tail, which served as anatomical regions of interest for all E-field metric extraction.

Subject-specific finite element head models were constructed in Sim4Life (v9.2, ZMT Zurich MedTech AG, Switzerland). Each participant’s T1-weighted image was resampled to 0.5 mm isotropic resolution using B-spline interpolation and segmented into 40 tissue types using Sim4Life’s AI-based deep learning pipeline (ImageML HeadModelGeneration, head40 model), including skin, fat, muscle, galea, compact and spongy bone, dura, cerebrospinal fluid, cortical grey matter, white matter, cerebellum, brainstem, subcortical structures, glands, and more. Subject-specific anisotropic white matter conductivity was incorporated by reconstructing diffusion tensors from each participant’s DWI data using the TensorModel implementation in dipy, with tensor components reordered to Sim4Life convention (XX, YY, ZZ, XY, YZ, ZX) prior to import. The preprocessed DWI data were coregistered to the T1-weighted image using a rigid-body transformation (6 degrees of freedom, Mattes mutual information similarity metric) implemented in SimpleITK. All remaining tissue conductivity values were assigned according to the low-frequency conductivity section of the IT’IS Tissue Properties Database v4.1^48^.

TIS electrode positions individually recorded during each sleep session using the Brainsight neuronavigation system (Rogue Research Inc., Montreal, Canada) with Polaris Vicra infrared optical tracking camera (Northern Digital Inc., Canada) and imported into each participant’s head model as custom cylindrical electrodes (radius = 7.5 mm), aligned to the head surface to ensure gap-less contact, matching the physical 1.5 cm diameter TENS electrodes used experimentally. Quasi-static finite element simulations were performed independently for each TIS electrode pair (pair 1: e1–e2; pair 2: e3–e4) using Sim4Life’s ElectroQsOhmicSimulation solver at 2000 Hz, with Dirichlet boundary conditions of +1.0 V and −1.0 V applied to the anode and cathode respectively, and remaining electrodes treated as perfect electric conductors. All fields were subsequently normalized to 1 mA total current. The computational grid was structured (voxel-based) and defined with a maximum step of 1.0 mm and electrode resolution of 0.1 mm.

The TIS modulation envelope distribution was computed using the maximum amplitude modulated (MAM) formula from Grossman et al. (2017)^18^ and scaled according to the experimentally applied currents (1 mA and 3 mA). Three TI exposure metrics were extracted for the left hippocampal head, body, and tail to characterise the spatial specificity of hippocampal targeting. Field strength was defined as the median TI MAM across voxels within the ROI. Selectivity was computed as the (power) ratio of the spatiotemporally - averaged square MAM within the target region and the same across the remainder of the brain, providing a dimensionless index of targeting focality. where values above 1.0 indicate preferential targeting of the ROI. Collateral exposure was defined as the percentage of non-target brain voxels receiving field strength exceeding the 90th percentile of the target field distribution, quantifying the extent of off-target exposure relative to stimulation intensity at the intended target.

### EEG

We recorded electroencephalography using an ActiChamp Plus amplifier system with 64 scalp active electrodes (actiCAP slim, Brain Products GmbH, Munich, Germany) plus auxiliary channels for electrooculography and electromyography (Brain Products GmbH, Munich, Germany). Scalp electrodes were filled with electrolyte gel (V20 Electrolyte Creme; Easycap GmbH, Germany) and impedances reduced to below 25 kΩ for all channels. Electrooculography was recorded using two electrodes placed 1 cm lateral and 1 cm superior to the outer canthi, referenced to contralateral mastoids. Electromyography was recorded from a chin electrode using low-impedance electrode cream (Lic2, CNSAC, Germany), with impedance below 50 kΩ at the start of the recording. Data were acquired using Lab Streaming Layer (LSL) on a Windows-based PC with three monitors displaying: (1) BrainVision LSL Viewer for real-time EEG visualisation, (2) stimulus presentation, and (3) LSL Recorder interface for data recording. During the nap, EEG was sampled at 10,000 Hz to capture high-frequency carrier signals and enable offline stimulation artefact removal. LSL Recorder captured four synchronised streams: actiCHamp EEG, actiCHamp markers, TI output from the NI-DAQ device, and PsychoPy markers containing event timing and trial outcomes.

### EEG Preprocessing

EEG data was preprocessed using EEGLAB toolbox (version 2021.1)^49^ in MATLAB 2024a (Mathworks, Natick, MA, USA). Filters were applied sequentially using EEGLAB’s pop_eegfiltnew function: high-pass at 0.5 Hz, low-pass at 30 Hz, and notch filter at 50 Hz (bandwidth 49-51 Hz). Data were then resampled to 256 Hz using pop_resample. Bad channels identified through manual inspection were removed and interpolated using spherical spline interpolation. Data were re-referenced to the average across all EEG channels, and epochs containing movement or other artefacts were marked and excluded from further analysis. Visual inspection before and after preprocessing revealed clean slow oscillations with characteristic sharp down-state transitions and spindles with typical waxing-and-waning amplitude envelopes, with no visible high-frequency temporal interference contamination or distortion of waveform morphology.

### Sleep Scoring

Sleep stages were scored offline according to American Academy of Sleep Medicine (AASM) guidelines^50^ using standard 30 s epochs in Domino software (Somnomedics, Germany). Scoring was performed blind to experimental conditions and trial timing to prevent bias. We computed bilateral EEG derivations for visualisation and scoring: F4-A1, C4-A1, P4-A1, O2-A1, F3-A2, C3-A2, P3-A2, and O1-A2, referenced to contralateral mastoids (A1 = FT9, A2 = FT10), providing comprehensive coverage of frontal, central, parietal, and occipital regions. EOG and EMG channels were displayed alongside EEG derivations for complete polysomnographic assessment. Each 30 s epoch was assigned to one sleep stage (Wake, N1, N2, N3, or REM) based on standard AASM criteria including EEG frequency content, amplitude, sleep spindles, K-complexes, eye movements, and muscle tone. Sleep architecture metrics computed for each participant included total sleep time, sleep efficiency, wake after sleep onset, and percentages and durations of each sleep stage.

### Sleep spindle and SO event detection

Slow oscillations and spindle events were detected in N2 and N3 sleep across all 64 EEG channels using the YASA toolbox^51^. For spindles, separate detection runs were conducted for fast (12-16 Hz) and slow (10-12 Hz) spindles. Spindles were identified using YASA’s default multi-threshold algorithm, which applies criteria based on sigma power, waveform correlation (Morlet wavelet), and root-mean-square amplitude, with duration constraints of 0.3-3 s. For slow wave detection, data were band-pass filtered at 0.5-2 Hz using YASA’s default finite impulse response filter, and slow wave events identified using the following amplitude thresholds: negative half-wave amplitude between 40-200 μV, positive half-wave amplitude between 15-150 μV, and peak-to-peak amplitude between 35-350 μV.

### SO-spindle coupling

For SO-spindle coupling, spindles were those already detected as described above. SOs for coupling analyses were detected for each channel using a custom MATLAB script to allow spindle-locked detection. EEG data were bandpass filtered to 0.5-2.0 Hz using a 4th-order zero-phase Butterworth filter. SO events were identified as half-waves spanning consecutive zero-crossings with durations of 0.8-2.0 s, with troughs required to occur within 0.25-3.75 s post-cue. Amplitude thresholds were calculated per participant and per channel as mean trough-to-peak amplitude of candidate events multiplied by 1.25, ensuring that only robust slow oscillation events were retained^23^. SO-spindle co-occurrence rate was computed as the mean number of fast spindle peaks occurring within ±1.5 seconds of a SO trough detected on same channel per valid N2/N3 trial, expressed as a percentage, computed independently per channel. Coupling strength was indexed as the mean vector length (MVL) of the SO phase distribution at fast spindle peaks, computed as the mean resultant length of the phase angles^52^. The instantaneous SO phase was computed via the Hilbert transform applied to the 0.5-2.0 Hz filtered signal, and the SO phase at each spindle peak occurring within 0.25–3.75 s post-cue was extracted per channel and trial. Group-level MVL topographical maps were computed by averaging coupling strength across participants per channel per condition, motivating the definition of a combined region of interest comprising 27 frontal, frontocentral, central, and centroparietal channels used for all subsequent phase analyses (Fpz, F1, F2, F3, F4, F5, F6, Fz, FC1, FC2, FC3, FC4, FC5, FC6, FCz, C1, C2, C3, C4, C5, C6, Cz, CP1, CPz, CP2, Fp1, Fp2). The preferred SO phase of spindle occurrence was computed for each participant as the circular mean of preferred phase angles across ROI channels.

### Data analysis

#### Behavioural data analysis

Behavioural analyses focused on characterising memory performance before and after sleep across three outcome measures: word recognition (correct identification of the word as old regardless of image category), associative memory (correct identification of both the word and its paired image category), and memory fidelity (the richness of the verbal description of the associated image). We focused on forgetting as the outcome of interest, as memory declines naturally over time. For word recognition and associative memory, forgetting was quantified at the item level among items correctly recalled pre-nap, coded 1 if the item was still correctly recalled post-nap (retained) and 0 if it was no longer correctly recalled (forgotten). Forgetting rates are reported throughout as 1 − retention probability. Memory fidelity was scored on a four-point scale: 0 (no recall), 1 (related but confused), 2 (correct general category), and 3 (accurate detailed description). To capture graded loss of detail, fidelity degradation was quantified at the item level as the pre-nap minus post-nap score, with higher values indicating greater loss of detail across the nap. One participant was excluded from fidelity analyses owing to an audio recording failure that prevented transcription, yielding a final fidelity sample of 27 participants.

Word recognition accuracy, associative memory accuracy, and mean memory fidelity were each calculated per participant in both pre- and post-nap sessions and compared using paired-samples t-tests

To investigate whether post-nap memory differed across experimental conditions, word recognition and associative memory forgetting were each modelled using separate GLMMs with binomial error distribution and logit link function. Each model included condition as a four-level categorical fixed effect (TIS+Cue, TIS+Cue delayed, Cue, No-Cue) and a random intercept for participant. Omnibus condition effects were assessed using Type II Wald chi-square tests. Memory fidelity was instead modelled using a linear mixed-effects model (LMM) with the same fixed and random effects structure, and the omnibus condition effect assessed by F-test. Post-hoc pairwise contrasts were computed using estimated marginal means.

All behavioural analyses were conducted in R version 4.5.0. Generalised linear mixed-effects models were implemented using the lme4 and lmerTest packages with Wald tests for fixed-effect estimates. Model diagnostics were performed using the DHARMa package, estimated marginal means were computed using the emmeans package, and model fit statistics were obtained using the performance package. Effects were considered statistically significant at p ≤ 0.05.

#### Spindle amplitude analysis

To examine condition effects on spindle amplitude, a channel-wise linear mixed-effects model (LME) was fitted per channel with, spindle amplitude as the dependent variable, condition (TIS+Cue, TIS+Cue delayed, Cue) as a fixed effect, trial number as a covariate, and participant as a random intercept (spindle amplitude ∼ condition + trial number + (1|participant)). An omnibus F-statistic for the condition effect was extracted from each channel-wise model using analysis of variance. To correct for multiple comparisons across channels, a cluster-based permutation test was applied. Significant channels (uncorrected p < 0.05) were grouped into spatial clusters using a triangulation-based neighbour structure derived from the actiCAP 64-channel standard layout, with a minimum cluster size of two channels. Observed cluster masses were defined as the sum of F-statistics within each cluster. A null distribution of maximum cluster masses was generated by performing 1,000 sign-flip permutations on participant-level condition means, with a fixed random seed to ensure reproducibility. Clusters were considered significant if their observed mass exceeded the 95th percentile of the null distribution (cluster-corrected p < 0.05). For significant clusters, mean spindle amplitude was averaged across cluster channels per participant and condition, and pairwise comparisons between conditions were conducted using LMEs with participant as a random intercept. Throughout all topographic maps, white dots indicate channels with significant effects and grey dots indicate non-significant channels. This procedure was run separately for fast and slow spindles.

#### SO amplitude analysis

To examine condition effects on slow oscillation amplitude, a channel-wise LME was fitted per channel with SO amplitude as the dependent variable, condition (TIS+Cue, TIS+Cue delayed, Cue) as a fixed effect, trial number as a covariate, and participant as a random intercept (SO amplitude ∼ condition + trial number + (1|participant)). An omnibus F-statistic for the condition effect was extracted from each channel-wise model using analysis of variance. The same cluster-based permutation approach described previously was applied to correct for multiple comparisons across channels. For significant clusters, mean SO amplitude was averaged across cluster channels per participant and condition, and pairwise comparisons between conditions were conducted using LMEs with participant as a random intercept.

#### Spindle density analysis

To examine condition effects on spindle density, a channel-wise GLMM with a Gamma distribution and was fitted per channel with spindle density as the dependent variable, condition (TIS+Cue, TIS+Cue delayed, Cue) as a fixed effect, and participant as a random intercept (Spindle density ∼ condition + (1 | participant), family = Gamma (link = “log”)). A Gamma distribution was used for all density analysis to account for the right-skewed nature of density data, in which values cluster near zero. An omnibus F-statistic for the condition effect was extracted from each channel-wise model using analysis of variance. The same cluster-based permutation approach described previously was applied to correct for multiple comparisons across channels. For significant clusters, mean spindle density was averaged across cluster channels per participant and condition, and pairwise comparisons between conditions were conducted using LMEs with participant as a random intercept. This procedure was run separately for fast and slow spindles.

#### SO density analysis

To examine condition effects on slow oscillation density, a channel-wise GLMM with a Gamma distribution and was fitted per channel with SO density as the dependent variable, condition (TIS+Cue, TIS+Cue delayed, Cue) as a fixed effect, and participant as a random intercept (SO density ∼ condition + (1 | participant), family = Gamma (link = “log”)). An omnibus F-statistic for the condition effect was extracted from each channel-wise model using analysis of variance. The same cluster-based permutation approach described previously was applied to correct for multiple comparisons across channels. For significant clusters, mean SO density was averaged across cluster channels per participant and condition, and pairwise comparisons between conditions were conducted using LMEs with participant as a random intercept

#### SO-spindle coupling analysis

Coupling rate and MVL, were each analysed using channel-wise LMEs model (coupling rate or MVL ∼ condition + trial + (1|participant)) with condition (TIS+Cue, TIS+Cue delayed, Cue) as a fixed effect, trial number as a covariate, and participant as a random intercept. The same cluster-based permutation approach described previously was applied to correct for multiple comparisons across channels for both measures. Phase clustering significance was assessed using the Rayleigh test, and between-condition differences in preferred SO phase were assessed using the Watson-Williams test. Brain-behaviour associations between SO-spindle coupling rate and associative memory-based forgetting were examined using channel-wise Pearson correlations across all 64 electrodes, with cluster-based permutation correction applied separately per condition (1,000 permutations).

### Mediation Analysis

#### EEG feature extractions

All artefact-free stimulation epochs were retained irrespective of sleep stage; 88.0– 95.9% of analysed trials across conditions fell within N2/N3, with closely matched N2:N3 composition across conditions. Stimulation events (Cue, TIS+Cue, TIS+Cue delayed) were identified from EEG markers, and for each event (regardless of condition), we extracted a 0.8 s baseline (−1 to −0.2 s before sound stimulus; in TIS+Cue delayed, this overlaps with the preceding TIS period), and a 2.8 s post-stimulus EEG segment (from 0.2 to 3 s after sound stimulus). Each trial was linked to the corresponding post-nap associative memory accuracy by the actual word that was delivered, with the associative memory accuracy defined as the correctness of the image-category (face vs object) response following a correct Old/New judgement. Trials with an incorrect Old/New judgment were coded as associative memory failures (0), since recollection cannot be expressed for an item not recognised as old, and to preserve class balance in the binary outcome. Foil trials were discarded. Then, for each trial, we computed eight EEG features within two windows (baseline and post-stimulus) defined relative to sound onset. The features comprised four spectral measures (spindle-band power 12–16 Hz, spindle peak frequency, theta power 4–8 Hz, and the spectral slope), one envelope measure (sigma band burst amplitude), and three phase–amplitude coupling (PAC) measures: SO–spindle, delta–spindle, and their differences. For the periodic spectral features, the EEG was band-pass filtered (0.1–32 Hz) and a differentiation filter was then applied to attenuate the 1/f spectral component and better capture the periodic component44. The power spectral density (PSD) of each window was estimated by Thomson’s multitaper method (MATLAB pmtm; time-half-bandwidth product NW = 4, i.e. seven Slepian tapers; FFT length set to the next power of two of the segment) and normalised to total spectral power, so that band power is expressed as a fraction of the whole spectrum. Band power was the log-transformed sum of the normalized PSD across bins within a band (theta 4–8 Hz; spindle 12– 16 Hz), and spindle peak frequency was the bin of maximum normalized power within 12–16 Hz.

The spectral slope (or the aperiodic component of the EEG power spectrum) was evaluated using the FOOOF (fitting oscillations and one over f, https://fooof-tools.github.io/fooof/) toolbox^54^, with a Python v.3.10 environment. The spectral slope was estimated at the broadband 1-30 Hz, with a larger value indicating a steeper slope. The evaluation was done without a differentiation filter prior. Sigma burst amplitude was the mean of the spindle-band (12–16 Hz) analytic envelope, obtained by zero-phase Butterworth band-pass filtering and the Hilbert transform. PAC was quantified by the mean vector length (MVL)^55^. For a low-frequency phase series *φ*(*t*) and a high-frequency amplitude series *A*(*t*) of length *T*, MVL is the modulus of the mean complex composite signal, normalised by mean amplitude:

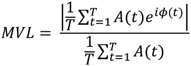

with spindle-band amplitude coupled to SO-band (0.5–2 Hz) or delta-band (2–4 Hz) phase. Band-pass filtering and the Hilbert transform were applied to the full 5 s segment before windowing, to avoid edge artefacts from filtering short epochs. We refer to this measure as PAC to distinguish it from the event-based coupling analysis above. Although, both use mean vector length, PAC is amplitude-weighted and computed continuously across all timepoints, rather than from the phase of SO relative to discrete detected spindle peaks.

Three additional cross-channel coupling features took the SO or delta phase from frontal channels and the spindle amplitude from central channels, averaged across all eligible frontal × central pairs (SO–spindle, delta–spindle, and their difference). All features were expressed as post-stimulus value minus baseline value, then averaged across channels within each region (Frontal, Central, left Temporal, right Temporal, Posterior, Occipital), yielding one trial-level value per feature per region. The Frontal region includes channels F1, F2, Fz, Fp1, Fp2, F3, F4, F5, F6, F7, F8, AFz, AF3, AF4, AF7 and AF8. The Central region includes channels Cz, C1, C2, C3, C4, C5, C6, FCz, FC1, FC2, FC3, FC4, FC5, FC6, CPz, CP1, CP2, CP3, CP4, CP5 and CP6. The left Temporal region includes channels FT7, FT9, T7, TP7 and TP9; the right Temporal region includes channels FT8, FT10, T8, TP8 and TP10. The Posterior region includes channels Pz, P1, P2, P3, P4, P5, P6, P7 and P8, and the Occipital region includes channels POz, PO3, PO4, PO7, PO8, Oz, O1 and O2.

#### Feature collinearity reduction

To mitigate strong collinearity among the 51 features (8 features × 6 regions plus 3 cross-channel coupling features), Spearman rank correlations were computed across all features over all retained trials and converted to a distance matrix (d = 1 − ∣ *r* ∣). Features were clustered by hierarchical agglomerative clustering with average linkage, and the dendrogram was cut at a distance of 0.5 (i.e. ∣ *r* ∣> 0.5, within a cluster). From each resulting cluster, we retained a single representative feature: the member whose absolute-correlation profile was closest (smallest Euclidean distance) to the cluster mean profile, with singleton clusters retained directly. This yielded 16 representative features, which were then used in all later analyses **(Supplementary Fig 11).** Because each mediated effect is estimated as the unique contribution of one feature conditional on all others, we verified that residual collinearity among the retained features was low: the strongest pairwise correlation was |r| = 0.56 and variance inflation factors in the joint model did not exceed 2.05 across contrasts (mean 1.35), well below conventional thresholds. Feature selection used no condition or outcome information.

#### Mediation analysis: Path-a (TIS effect on EEG features)

The treatment effect on each EEG feature (path a, X → M) was tested as a standalone analysis, separately for each of the three pairwise condition contrasts: TIS+Cue vs Cue, TIS+Cue delayed vs Cue, and TIS+Cue vs TIS+Cue delayed. For each feature and each contrast, the trial-level feature was z-scored and modelled by a linear mixed model with a fixed effect of condition and a participant random intercept, with degrees of freedom approximated by the Satterthwaite method (lme4/lmerTest),

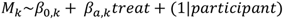

where *M_k_* is the z-scored value of feature *k* (the mediator), and *β_a_*_,*k*_ is its treatment effect; treat is a binary indicator coding the two conditions of the contrast (the first-named condition = 1, the reference condition = 0; e.g., for TIS+Cue vs Cue, TIS+Cue = 1 and Cue = 0). The coefficient *β_a_* and its p-value quantify the treatment effect on that feature. Trials were retained whenever the feature was defined (mediator-valid), independent of the behavioural outcome, and participants with fewer than 5 trials per condition were excluded from that contrast. Within each contrast, p-values were FDR-corrected (Benjamini–Hochberg) across features. Path-a results are reported in **Supplementary Fig 12** and **Supplementary Table 8.**

#### Mediation effect estimations

To estimate the indirect effect attributable to each EEG feature while accounting for their shared variance, we fitted a single joint mediation model per contrast, in which all selected features act as parallel mediators of associative memory accuracy. Each feature was z-scored across trials. The mediator models were the per-feature path-a models above. The outcome model was a single generalised linear mixed model (binomial logit) regressing associative memory accuracy on all mediators and the treatment simultaneously,

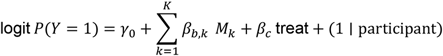

The mediators *M_k_* are the same z-scored features modelled in path a; each path-b coefficient *β_b_*_,*k*_ reflects the unique contribution of *M_k_* controlling for all other features, and the per-feature ACME combines the path-a model *M_k_* with this joint outcome model. The average causal mediation effect (ACME) for each feature was then obtained within this joint model by quasi-Bayesian Monte Carlo approximation^56^ (R package mediation, 1000 simulations, no bootstrap), pairing the feature’s mediator model with the joint outcome model. For each feature, we recorded the ACME (posterior mean, SD, and the normalised ACME defined as mean/SD), the path-b estimate and p-value, the average direct effect, and the proportion mediated. Models were fitted on the binary associative memory accuracy outcome with a participant random intercept; participants with fewer than 5 trials per condition were excluded per contrast, and joint-model singularity was recorded, but no models were singular. Only the TIS+Cue vs Cue contrast is reported in the main text; full per-contrast statistics are provided in **Supplementary Table 8**.

#### Surrogate testing for indirect effect significance evaluation

To evaluate the significance of the ACME per feature, we constructed an empirical null distribution by permuting the treatment label, as a non-parametric complement to the quasi-Bayesian interval. On each permutation, the condition label was shuffled within participant, preserving each participant’s per-condition trial counts, and the entire pipeline (per-feature path-a, joint path-b, and per-feature ACME) was refitted on the permuted data to yield one null ACME per feature (200 quasi-Bayesian simulations per refit). For each feature, the one-sided permutation p-value was the right-tailed exceedance probability of the observed ACME underthis null, 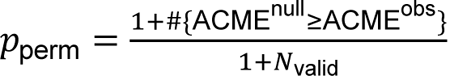, where *N*_valid_ is the number of permutations yielding a valid refit.

#### Mediation statistics summary

To assess whether the indirect effects across features showed a coordinated direction, we computed the normalised ACME (ratio of the ACME posterior mean to its SD across simulation draws), a sign-preserving, unit-free signal-to-noise summary of the indirect effect, for each feature. The distribution of normalised ACMEs across features was tested against zero with a two-sided one-sample t-test, separately for each contrast. The regional summary in **Fig. 5d** was obtained by averaging the normalised ACMEs across all features within each region and rendering each region as a fixed-radius bubble at a representative scalp location, colour-coded by its mean.

#### TI field metrics analysis

Differences in E-field strength, selectivity, and collateral stimulation across hippocampal subregions (HP-Head, HP-Body, HP-Tail) were assessed using linear mixed-effects models (LME) with subregion as a fixed effect and participant as a random intercept. Pairwise contrasts were conducted using estimated marginal means with FDR correction.

#### TI field metrics - forgetting analysis

To examine whether individual differences in hippocampal head (HP-Head) E-field strength, selectivity, and collateral exposure predicted associative memory forgetting, linear mixed-effects models (LMEs) were fitted for each metric (forgetting ∼ condition × metric + (1|participant)), with forgetting (%) as the dependent variable, condition, the E-field metric, and their interaction as fixed effects, and participant as a random intercept. Fixed effects were tested using Type II Wald F-tests with Kenward-Roger degrees of freedom. Each model was then repeated including each participant’s Cue-only forgetting rate as a covariate (forgetting ∼ condition × metric + CueOnlyForgetting + (1|participant)), to test whether effects were specific to the field metric rather than reflecting a general tendency for participants who forgot more in one condition to also forget more in the other. Condition-specific slopes for each metric were estimated using estimated marginal trends (emtrends).

### Subjective Measures

#### Subjective sleepiness

Before and after the afternoon nap, we assessed subjective sleepiness using the Karolinska Sleepiness Scale (KSS)^57^. The KSS is a validated 9-point scale ranging from 1 (very alert) to 9 (extremely sleepy, fighting sleep). Participants completed the scale electronically via Microsoft Forms immediately before the nap and again immediately upon awakening. Participants felt significantly less sleepy and more alert following the nap. Pre- and post-nap statistical comparisons are provided in **Supplementary Table 11**.

### Sleep quality and restfulness

Upon awakening from the nap, participants rated their sleep quality on a 6-point scale from 1 (very poor) to 6 (excellent) and post-sleep refreshment on a 5-point scale from 1 (not at all rested) to 5 (extremely rested). Ratings are provided in **Supplementary Table 12.**

#### Dream experience

We assessed dream experience with yes/no/maybe response to whether participants dreamt during the nap. If affirmed, participants rated on a four-point scale (not at all, at times, mostly, exclusively) whether dreams related to task content (faces/objects/sounds), related to the experimental context (being in the lab), were unrelated to task or lab, were pleasant, or were unpleasant. This provided insight into whether memory reactivation during sleep influenced dream content. Out of the 28 participants, 7 reported dreaming during the nap, and 9 were unsure if they dreamed. Among the 16 participants who reported dreaming or unsure (yes or maybe), 11 indicated their dreams were not at all related to the task content, 3 indicated their dreams were related at times, and 2 did not respond to this question.

#### Awareness of Auditory Cues

Awareness of auditory cues was assessed by asking participants whether they had heard any sounds while sleeping or attempting to sleep (yes / no / unsure). Those endorsing awareness provided free-text descriptions of the sounds perceived, their approximate timing within the nap, and their emotional response. Six out of 28 participants (21%) reported hearing auditory memory cues during the nap. Emotional responses were predominantly neutral or indifferent (n = 3), with some participants reporting annoyance or irritation (n = 2), or being woken by the sounds (n = 1). Similar post-sleep self-report measures have been used in prior targeted memory reactivation studies, in which a subset of participants reported hearing task-related sounds during sleep^58^. However, given our within-subject design, it is implausible that cue awareness differed systematically across conditions, and therefore unlikely to explain the timing-dependent effects observed.

## Code and Data Availability

The code for the memory task is available on https://github.com/nsnlab/MYSTI_Memory_Task. Data analysis scripts are available on https://github.com/nsnlab/TIS_Sleep_Memory. MRI and EEG data are available in https://zenodo.org/records/22210654.

## ACKNOWLEDGEMENTS

P.O, U.B and I.R.V. are supported by the Biotechnology and Biological Sciences Research Council (BB/Y011856/1). P.O. and T.R. received doctoral fellowships from the University of Surrey. VJ was supported by the Swiss National Science Foundation (P2EZP3_199918, P500PB_217827), the UK Dementia Research Institute (UKDRI-7206), Care Research and Technology Centre at Imperial College, London, and the University of Surrey, Guildford and Wellcome (301007/Z/23/Z). DJD is supported by the UK Dementia Research Institute (UKDRI-7206), Care Research and Technology Centre at Imperial College, London, and the University of Surrey, Guildford, and by the NIHR Oxford Health Biomedical Research Centre (NIHR 203316).

## AUTHOR CONTRIBUTIONS

CRediT: PO: Conceptualization, Methodology, Formal analysis, Investigation, Data Curation, Visualization, Writing - Original Draft, Writing - Review & Editing; JL: Formal analysis, Visualization, Writing - Original Draft, Review & Editing; TR: Software, Investigation, Formal analysis; KA: Methodology; DLJ: Investigation; MS: Software; EN: Resources, Software; NK: Resources; NG: Methodology, Writing - Review & Editing; DJD: Writing - Review & Editing, Funding acquisition; RCK: Supervision, Writing - Review & Editing; UB: Resources, Writing - Review & Editing, Funding acquisition; VJ: Formal analysis, Supervision, Writing - Review & Editing; IRV: Conceptualization, Methodology, Resources, Supervision, Project administration, Funding acquisition, Writing - Original Draft, Review & Editing.

## REFERENCES

1. Born, J. & Wilhelm, I. System consolidation of memory during sleep. Psychol. Res. 76, 192–203 (2012).

2. Diekelmann, S. & Born, J. The memory function of sleep. Nat. Rev. Neurosci. 11, 114–126 (2010).

3. Klinzing, J. G., Niethard, N. & Born, J. Mechanisms of systems memory consolidation during sleep. Nat. Neurosci. 22, 1598–1610 (2019).

4. Rasch, B. & Born, J. About Sleep’s Role in Memory. Physiol Rev 93, 681–766 (2013).

5. Axmacher, N., Elger, C. E. & Fell, J. Ripples in the medial temporal lobe are relevant for human memory consolidation. Brain 131, 1806–1817 (2008).

6. Staresina, B. P. et al. Hierarchical nesting of slow oscillations, spindles and ripples in the human hippocampus during sleep. Nat. Neurosci. 18, 1679–1686 (2015).

7. Staba, R. J., Wilson, C. L., Bragin, A. & Fried, I. Quantitative analysis of high-frequency oscillations (80-500 Hz) recorded in human epileptic hippocampus and entorhinal cortex. J. Neurophysiol. 88, (2002).

8. Mölle, M., Bergmann, T. O., Marshall, L. & Born, J. Fast and slow spindles during the sleep slow oscillation: Disparate coalescence and engagement in memory processing. Sleep 34, 1411–1421 (2011).

9. Girardeau, G., Benchenane, K., Wiener, S. I., Buzsáki, G. & Zugaro, M. B. Selective suppression of hippocampal ripples impairs spatial memory. Nat. Neurosci. 12, 1222–1223 (2009).

10. Fernández-Ruiz, A. et al. Long-duration hippocampal sharp wave ripples improve memory. Science (1979). 364, (2019).

11. Muehlroth, B. E. et al. Precise Slow Oscillation–Spindle Coupling Promotes Memory Consolidation in Younger and Older Adults. Sci. Rep. 9, (2019).

12. Ng, T., Noh, E. & Spencer, R. M. Bayesian meta-analysis reveals the mechanistic role of slow oscillation-spindle coupling in sleep-dependent memory consolidation. Elife 13, (2025).

13. Clemens, Z. et al. Fine-tuned coupling between human parahippocampal ripples and sleep spindles. European Journal of Neuroscience 33, (2011).

14. Latchoumane, C. F. V., Ngo, H. V. V., Born, J. & Shin, H. S. Thalamic Spindles Promote Memory Formation during Sleep through Triple Phase-Locking of Cortical, Thalamic, and Hippocampal Rhythms. Neuron 95, 424–435.e6 (2017).

15. Maingret, N., Girardeau, G., Todorova, R., Goutierre, M. & Zugaro, M. Hippocampo-cortical coupling mediates memory consolidation during sleep. Nat. Neurosci. 19, 959–964 (2016).

16. Siapas, A. G. & Wilson, M. A. Coordinated interactions between hippocampal ripples and cortical spindles during slow-wave sleep. Neuron 21, (1998).

17. Geva-Sagiv, M. et al. Augmenting hippocampal–prefrontal neuronal synchrony during sleep enhances memory consolidation in humans. Nat. Neurosci. 26, 1100–1110 (2023).

18. Grossman, N. et al. Noninvasive Deep Brain Stimulation via Temporally Interfering Electric Fields. Cell 169, 1029–1041.e16 (2017).

19. Violante, I. R. et al. Non-invasive temporal interference electrical stimulation of the human hippocampus. Nat. Neurosci. 26, 1994–2004 (2023).

20. Missey, F. et al. Temporal Interference Stimulation Can Enhance or Disrupt Human Memory Encoding as a Function of Brain Location and Frequency. (2025) doi:10.1101/2025.09.22.677714.

21. Hu, X., Cheng, L. Y., Chiu, M. H. & Paller, K. A. Promoting memory consolidation during sleep: A meta-analysis of targeted memory reactivation. Psychol. Bull. 146, 218–244 (2020).

22. Oudiette, D. & Paller, K. A. Upgrading the sleeping brain with targeted memory reactivation. Trends Cogn. Sci. 17, 142–149 (2013).

23. Cairney, S. A., Guttesen, A. á. V., El Marj, N. & Staresina, B. P. Memory Consolidation Is Linked to Spindle-Mediated Information Processing during Sleep. Current Biology 28, 948–954.e4 (2018).

24. Fuentemilla, L. et al. Hippocampus-dependent strengthening of targeted memories via reactivation during sleep in humans. Current Biology 23, (2013).

25. Yonelinas, A., Hawkins, C., Abovian, A. & Aly, M. The role of recollection, familiarity, and the hippocampus in episodic and working memory. Neuropsychologia 193, (2024).

26. Yonelinas, A. P. The nature of recollection and familiarity: A review of 30 years of research. J. Mem. Lang. 46, (2002).

27. Borders, A. A., Aly, M., Parks, C. M. & Yonelinas, A. P. The hippocampus is particularly important for building associations across stimulus domains. Neuropsychologia 99, (2017).

28. Richter, F. R., Cooper, R. A., Bays, P. M. & Simons, J. S. Distinct neural mechanisms underlie the success, precision, and vividness of episodic memory. Elife 5, (2016).

29. Nicolas, J. et al. Unraveling the neurophysiological correlates of phase-specific enhancement of motor memory consolidation via slow-wave closed-loop targeted memory reactivation. Nature Communications 16, (2025).

30. Ngo, H. V. V. & Staresina, B. P. Shaping overnight consolidation via slow-oscillation closed-loop targeted memory reactivation. Proc. Natl. Acad. Sci. U. S. A. 119, (2022).

31. Clemens, Z. et al. Temporal coupling of parahippocampal ripples, sleep spindles and slow oscillations in humans. Brain 130, 2868–2878 (2007).

32. Fehér, K. D. et al. Shaping the slow waves of sleep: A systematic and integrative review of sleep slow wave modulation in humans using non-invasive brain stimulation. Sleep Med. Rev. 58, (2021).

33. Schaeffer, E. L. et al. Enhancement of Sleep Slow Wave Activity using Transcranial Electrical Stimulation with Temporal Interference. Preprint at 10.1101/2025.08.11.25333452 (2025).

34. Missey, F. et al. Non-invasive temporal interference stimulation of the hippocampus suppresses epileptic biomarkers in patients with Epilepsy: biophysical differences between kilohertz and amplitude modulated stimulation. Brain Stimul. 19, (2026).

35. Helfrich, R. F., Mander, B. A., Jagust, W. J., Knight, R. T. & Walker, M. P. Old Brains Come Uncoupled in Sleep: Slow Wave-Spindle Synchrony, Brain Atrophy, and Forgetting. Neuron 97, 221–230.e4 (2018).

36. Bar, E. et al. Local Targeted Memory Reactivation in Human Sleep. Current Biology 30, (2020).

37. Schönauer, M. et al. Decoding material-specific memory reprocessing during sleep in humans. Nat. Commun. 8, (2017).

38. Dimitrov, T., He, M., Stickgold, R. & Prerau, M. J. Sleep spindles comprise a subset of a broader class of electroencephalogram events. Sleep 44, (2021).

39. Kasten, F. H., Duecker, K., Maack, M. C., Meiser, A. & Herrmann, C. S. Integrating electric field modeling and neuroimaging to explain inter-individual variability of tACS effects. Nat. Commun. 10, (2019).

40. Demchenko, I. et al. Human applications of transcranial temporal interference stimulation: A systematic review. Brain Stimul. 18, (2025).

41. Nissim, N. R. et al. Efficacy of Transcranial Alternating Current Stimulation in the Enhancement of Working Memory Performance in Healthy Adults: A Systematic Meta-Analysis. Neuromodulation 26, 728–737 (2023).

42. Creery, J. D., Oudiette, D., Antony, J. W. & Paller, K. A. Targeted memory reactivation during sleep depends on prior learning. Sleep 38, 755–763 (2015).

43. Antony, J. W., Gobel, E. W., O’Hare, J. K., Reber, P. J. & Paller, K. A. Cued memory reactivation during sleep influences skill learning. Nat. Neurosci. 15, 1114–1116 (2012).

44. Batterink, L. J., Westerberg, C. E. & Paller, K. A. Vocabulary learning benefits from REM after slow-wave sleep. Neurobiol. Learn. Mem. 144, 102–113 (2017).

45. Schreiner, T., Petzka, M., Staudigl, T. & Staresina, B. P. Endogenous memory reactivation during sleep in humans is clocked by slow oscillation-spindle complexes. Nat. Commun. 12, (2021).

46. Rasch, B., Büchel, C., Gais, S. & Born, J. Odor cues during slow-wave sleep prompt declarative memory consolidation. Science (1979). 315, 1426–1429 (2007).

47. Rudoy, J. D., Voss, J. L., Westerberg, C. E. & Paller, K. A. Strengthening individual memories by reactivating them during sleep. Science (1979). 326, 1079 (2009).

48. Hasgall, P., et al. IT’IS Database for thermal and electromagnetic parameters of biological tissues. Version 4.1, Feb 22 (2022).

49. Delorme, A. & Makeig, S. EEGLAB: An open source toolbox for analysis of single-trial EEG dynamics including independent component analysis. J. Neurosci. Methods 134, 9–21 (2004).

50. Berry, R. B., et al. AASM | Scoring Manual Version 2.4 The AASM Manual for the Scoring of Sleep and Associated Events RULES, TERMINOLOGY AND TECHNICAL SPECIFICATIONS VERSION 2.4. www.aasmnet.org. (2017).

51. Vallat, R. & Walker, M. P. An open-source, high-performance tool for automated sleep staging. Elife 10, 70092 (2021).

52. Berens, P. CircStat: A MATLAB Toolbox for Circular Statistics. J. Stat. Softw. 31, (2009).

53. Demanuele, C., James, C. J. & Sonuga-Barke, E. J. S. Distinguishing low frequency oscillations within the 1/f spectral behaviour of electromagnetic brain signals. Behavioral and Brain Functions 3, (2007).

54. Donoghue, T. et al. Parameterizing neural power spectra into periodic and aperiodic components. Nat. Neurosci. 23, (2020).

55. Canolty, R. T. & Knight, R. T. The functional role of cross-frequency coupling. Trends Cogn. Sci. 14, (2010).

56. Imai, K., Keele, L. & Tingley, D. A General Approach to Causal Mediation Analysis. Psychol. Methods 15, (2010).

57. Akerstedt, T. SUBJECTIVE AND OBJECTIVE SLEEPINESS IN THE ACTIVE INDIVIDUAL. vol. 52 (1990).

58. Schechtman, E., Heilberg, J. & Paller, K. A. Memory consolidation during sleep involves context reinstatement in humans. Cell Rep. 42, (2023).

