## Supplementary Material for "Hippocampal stimulation timed to memory reactivation shapes human sleep oscillatory dynamics and consolidation"

### Supplemental Information

##### Table S1 | TIS electric field metrics across left hippocampal subregions.

Mean ± SD of field strength, selectivity, and collateral exposure per subregion. Omnibus one-way repeated-measures ANOVA - Strength: $F(2,54) = 325.3$, $p < 2.2 \times 10^{-31}$, $\eta^2 = 0.92$; Selectivity: $F(2,54) = 569.5$, $p < 5.1 \times 10^{-37}$, $\eta^2 = 0.96$; Collateral: $F(2,54) = 642.2$, $p < 2.3 \times 10^{-38}$, $\eta^2 = 0.96$.

| Subregion | Field strength (V/m) Mean ± SD | Selectivity Mean ± SD | Collateral (%) Mean ± SD |
|---|---|---|---|
| **HP-Head** | 0.360 ± 0.061 | 1.96 ± 0.266 | 7.91 ± 2.16 |
| **HP-Body** | 0.279 ± 0.045 | 1.19 ± 0.167 | 18.6 ± 3.44 |
| **HP-Tail** | 0.190 ± 0.023 | 0.567 ± 0.112 | 36.3 ± 6.17 |

Pairwise FDR-corrected contrasts (Benjamini–Hochberg). All ps < .0001. SE = standard error. Selectivity = ratio of hippocampal to whole-brain mean envelope modulation; Collateral = percentage of brain volume receiving ≥ 50% of hippocampal field strength.

| Contrast | Estimate | SE | t ratio | p value (FDR) |
|---|---|---|---|---|
| **Strength (V/m)** | | | | |
| HP-Head – HP-Body | 0.080 | 0.007 | 12.03 | < .0001 |
| HP-Head – HP-Tail | 0.170 | 0.007 | 25.49 | < .0001 |
| HP-Body – HP-Tail | 0.090 | 0.007 | 13.46 | < .0001 |
| **Selectivity** | | | | |
| HP-Head – HP-Body | 0.768 | 0.041 | 18.59 | < .0001 |
| HP-Head – HP-Tail | 1.392 | 0.041 | 33.69 | < .0001 |
| HP-Body – HP-Tail | 0.624 | 0.041 | 15.10 | < .0001 |
| **Collateral (%)** | | | | |
| HP-Head – HP-Body | −8.70 | 0.813 | −10.71 | < .0001 |
| HP-Head – HP-Tail | −28.42 | 0.813 | −34.97 | < .0001 |
| HP-Body – HP-Tail | −19.72 | 0.813 | −24.27 | < .0001 |

*HP-Head = left hippocampal head; HP-Body = left hippocampal body; HP-Tail = left hippocampal tail*

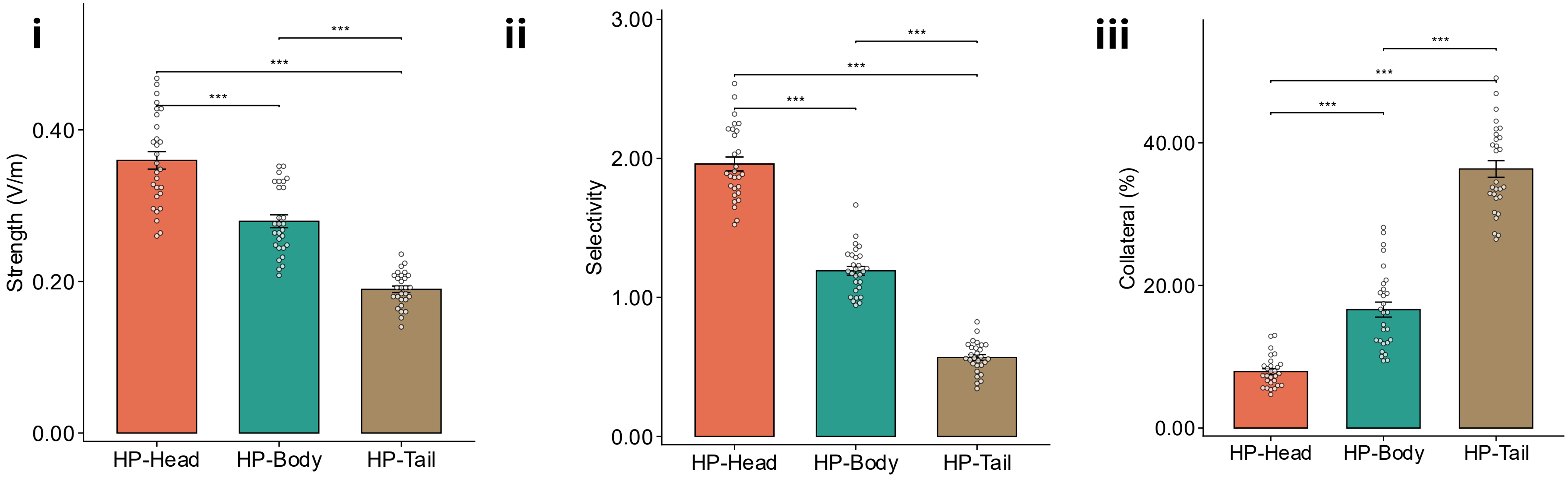


**Fig. S1 | TIS electric field metrics across left hippocampal subregions**

**(i)** TIS modulation strength (V/m), **(ii)** selectivity, and **(iii)** collateral exposure (%) across HP-Head, HP-Body, and HP-Tail (N=28). One-way repeated-measures ANOVA showed a significant anterior-posterior gradient for all three metrics: Strength: $F(2,54) = 325.3$, $p < 2.2\times10^{-31}$; Selectivity: $F(2,54) = 569.5$, $p < 5.1\times10^{-37}$; Collateral: $F(2,54) = 642.2$, $p < 2.3\times10^{-38}$. All pairwise contrasts (FDR-corrected) ps < .0001. ***p < .0001. Individual data points shown in the plots; bars show mean ± SEM

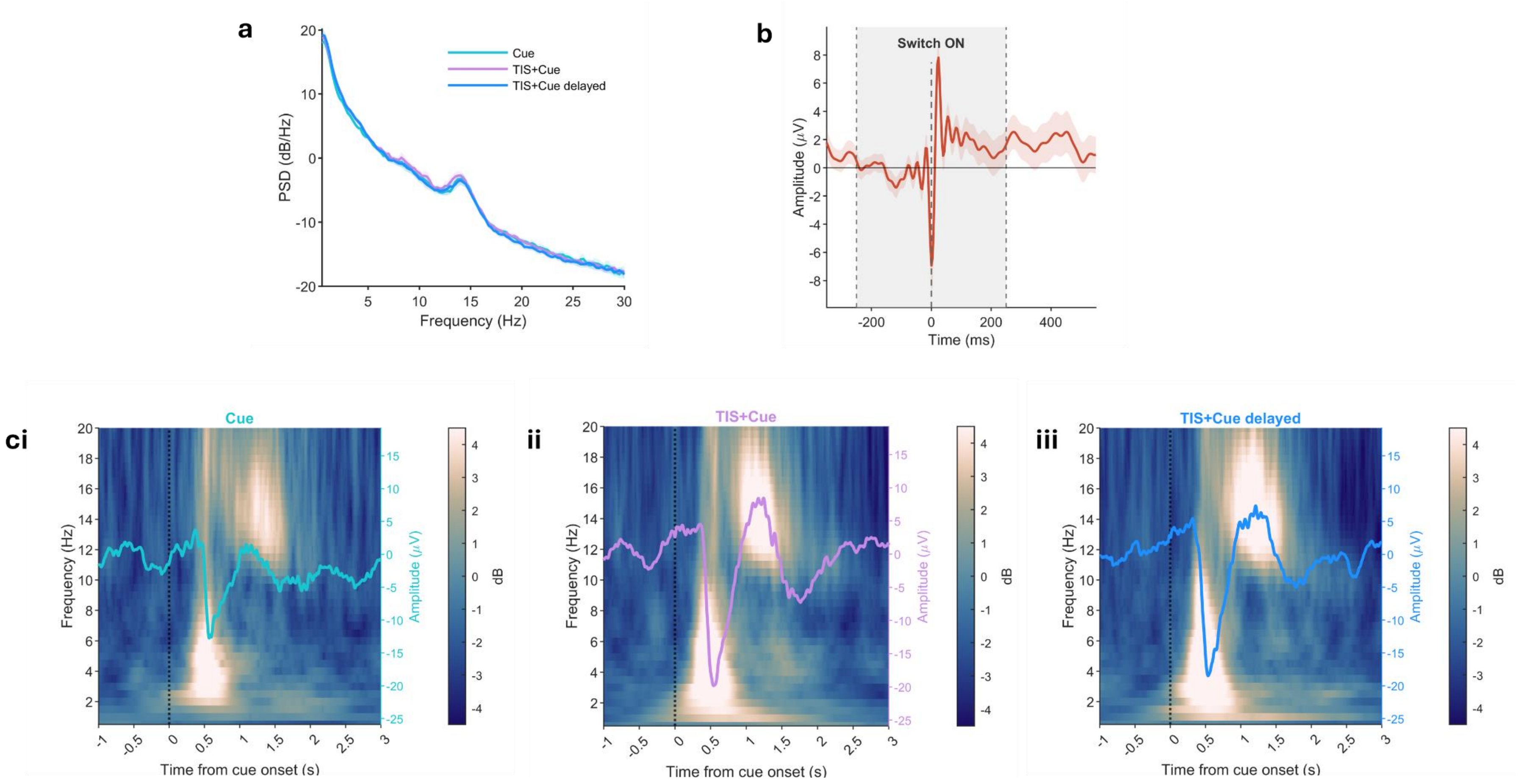


**Fig. S2 | EEG artifact control and signal quality during TIS**

**(a)** Power spectral density across conditions, showing preserved 1/f sleep EEG spectra with slow oscillation and spindle peaks. **(b)** Average EEG waveform time-locked to TIS frequency transition (t = 0, dashed line), showing a brief transient artifact (~100 ms) at the frequency switching. Grey shading indicates the ±250ms exclusion window applied to EEG event detection analyses. (c) Cue-locked time-frequency spectrograms showing power change (dB) relative to pre-cue baseline (−1.0 to −0.2 s before cue onset) for Cue **(i),** TIS+Cue **(ii),** and TIS+Cue delayed **(iii),** confirming that TIS delivery did not contaminate the cue-evoked EEG response. Overlaid waveforms show the average ERP amplitude (μV, right y-axis). Colour scale represents power change in dB.

**Table S2 | Sleep parameters (N = 28).**

Sleep architecture parameters recorded during the nap session.

| Parameter | Mean | SD |
|---|---|---|
| **Total sleep time (TST, min)** | 50.6 | 13.9 |
| **WASO (min)** | 9.3 | 9.5 |
| **Sleep efficiency (%)** | 84.4 | 16.1 |
| **N1 (% TST)** | 21.1 | 11.5 |
| **N2 (% TST)** | 66.8 | 14.4 |
| **N3 (% TST)** | 11.9 | 14.9 |
| **REM (% TST)** | 0.1 | 0.9 |

*WASO = waking time after sleep onset; TST = total sleep time.*

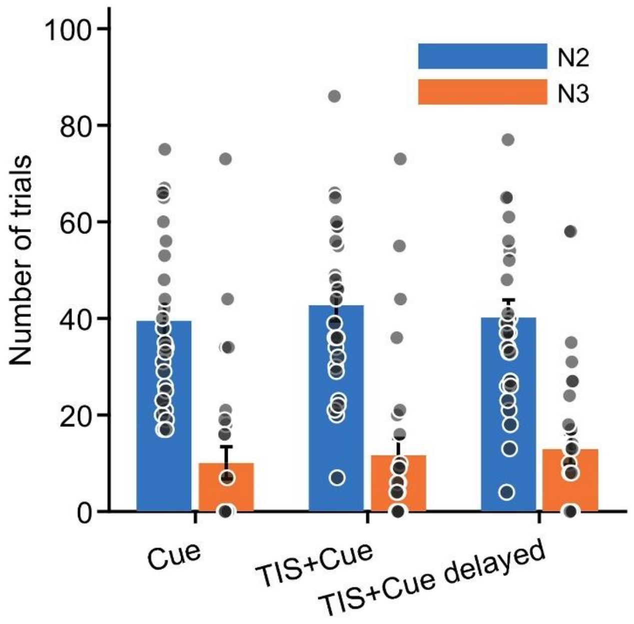


**Fig. S3 | Trials by sleep stage and condition**

Number of trials contributing to each condition (Cue, TIS+Cue, TIS+Cue delayed), by sleep stage (N2, N3). Main effect of sleep stage: $F(1,162) = 56.88$, $p < .001$; main effect of condition: $F(2,162) = 0.37$, $p = .690$; condition × sleep stage interaction: $F(2,162) = 0.24$, $p = .790$. Dots represent individual participants; bars show mean ± SEM, N=28.

##### Table S3 | Trial distribution across N2 and N3 sleep stages per condition.

Mean (± SD) number of trials per sleep stage across conditions. Significantly more trials occurred during N2 than N3 sleep (main effect of sleep stage: $F(1,162) = 56.88$, $p < .001$), reflecting the greater proportion of N2 in afternoon nap architecture. Conditions did not differ in trial distribution (main effect of condition: $F(2,162) = 0.37$, $p = .690$; condition × sleep stage interaction: $F(2,162) = 0.24$, $p = .790$).

| Condition | N2 trials (Mean ± SD) | N3 trials (Mean ± SD) |
|---|---|---|
| **TIS+Cue** | 42.7 ± 3.2 | 11.6 ± 3.5 |
| **TIS+Cue delayed** | 40.1 ± 3.7 | 12.9 ± 3.2 |
| **Cue** | 39.5 ± 3.3 | 10.1 ± 3.3 |

*SD, standard deviation*

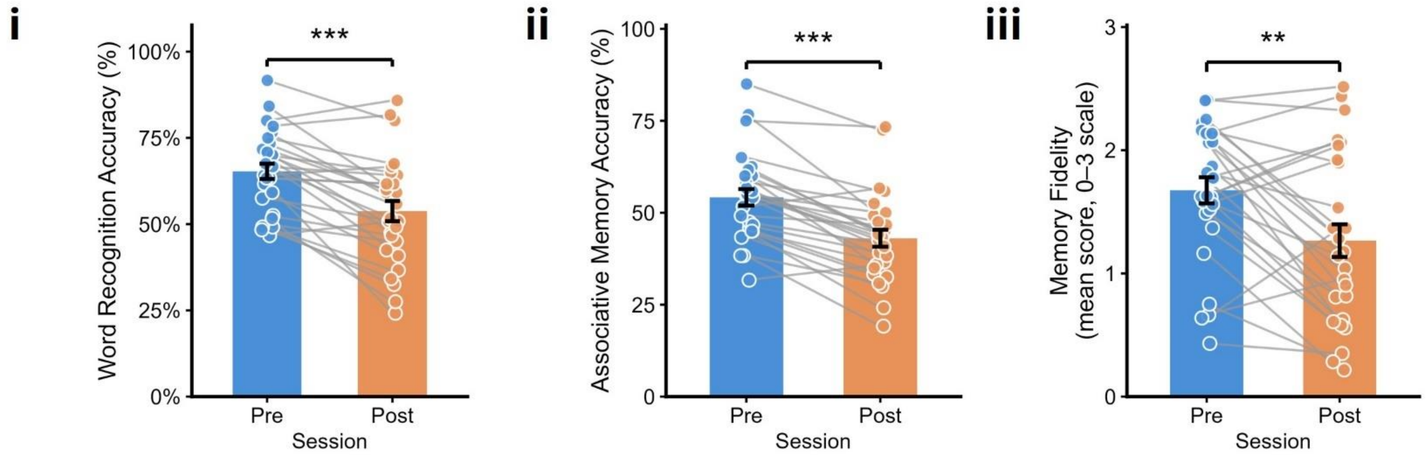


**Fig. S4 | Memory performance declines over time.**
**(i)** Word recognition accuracy, **(ii)** associative recall accuracy, and **(iii)** memory fidelity (0–3 scale, higher scores indicate better performance), before (Pre) and after (Post) the nap, for each participant (grey lines connect individual scores). Word recognition declined 11.5% from pre- to post-nap ($t(27) = -5.83$, $p < .001$, $d = 1.10$, N=28); associative memory declined 11.1% ($t(27) = -7.69$, $p < .001$, $d = 1.45$, N=28); memory fidelity declined from 1.67 to 1.27 ($t(26) = -3.49$, $p = .002$, $d = 0.67$, N=27). Bars show mean ± SEM; dots represent individual participants.

#### Table S4 | Associative memory GLMM - Estimated marginal means.

Estimated marginal means (forgetting probability) from associative memory. Overall condition effect: $\chi^2(3) = 10.66$, $p = 0.014$.

| Condition | Forgetting probability | SE | 95% CI lower | 95% CI upper |
|---|---|---|---|---|
| **TIS+Cue** | 0.059 | 0.015 | 0.036 | 0.096 |
| **TIS+Cue delayed** | 0.101 | 0.022 | 0.065 | 0.153 |
| **Cue** | 0.074 | 0.018 | 0.046 | 0.117 |
| **No-Cue** | 0.118 | 0.024 | 0.078 | 0.175 |

| Contrast | B | SE | z | p-value |
|---|---|---|---|---|
| **TIS+Cue - TIS+Cue delayed** | -0.590 | 0.267 | -2.209 | 0.027 |
| **TIS+Cue - Cue** | -0.250 | 0.277 | -0.901 | 0.368 |
| **TIS+Cue - No-Cue** | -0.765 | 0.257 | -2.977 | 0.003 |

| Contrast | B | SE | z | p-value |
|---|---|---|---|---|
| **TIS+Cue delayed - Cue** | 0.340 | 0.256 | 1.326 | 0.185 |
| **TIS+Cue delayed - No-Cue** | -0.175 | 0.234 | -0.748 | 0.455 |
| **Cue - No-Cue** | -0.515 | 0.246 | -2.091 | 0.037 |

*B = log odds estimate; SE = standard error.*

##### Table S5 | Fidelity GLMM - Estimated marginal means.

Estimated marginal means (pre – post scores on 0 – 3 scale) from the fidelity LMM. Overall condition effect: $F(3, 78) = 3.03$, $p = 0.034$.

| Condition | Mean index | 95% CI lower | 95% CI upper |
|---|---|---|---|
| **TIS+Cue** | 0.486 | 0.286 | 0.685 |
| **TIS+Cue delayed** | 0.782 | 0.583 | 0.982 |
| **Cue** | 0.676 | 0.477 | 0.876 |
| **No-Cue** | 0.718 | 0.519 | 0.918 |

| Contrast | B | t | p-value |
|---|---|---|---|
| **TIS+Cue - TIS+Cue delayed** | −0.297 | −2.861 | 0.005 |
| **TIS+Cue - Cue** | −0.191 | −1.838 | 0.070 |
| **TIS+Cue - No-Cue** | −0.233 | −2.242 | 0.028 |
| **TIS+Cue delayed - Cue** | 0.106 | 1.023 | 0.309 |
| **TIS+Cue delayed - No-Cue** | 0.064 | 0.619 | 0.538 |
| **Cue - No-Cue** | −0.042 | −0.405 | 0.687 |

*B = mean difference in forgetting index; SE = standard error.*

##### Table S6 | Linear mixed-effects models of hippocampal head E-field metrics on associative memory-based forgetting.

Type II Wald F-tests with Kenward-Roger degrees of freedom, without and with each participant's Cue-only forgetting rate as a covariate

| | Predictors | F (no covariate) | df (no covariate.) | p (no covariate.) | F (with covariate.) | df (with covariate.) | p (with covariate.) |
|---|---|---|---|---|---|---|---|
| **Strength** | Condition | 6.25 | 1,26 | 0.019 | 6.25 | 1,26 | 0.019 |
| | HP_Head_Strength | 5.61 | 1,26 | 0.026 | 2.20 | 1,25 | 0.150 |
| | CueOnlyForgetting | — | — | — | 6.35 | 1,25 | 0.019 |
| | Condition × Strength | 0.03 | 1,26 | 0.874 | 0.03 | 1,26 | 0.874 |
| **Selectivity** | Condition | 7.23 | 1,26 | 0.012 | 7.23 | 1,26 | 0.012 |
| | HP_Head_Strength | 0.57 | 1,26 | 0.456 | 0.80 | 1,25 | 0.378 |
| | CueOnlyForgetting | — | — | — | 10.37 | 1,25 | 0.003 |
| | Condition × Selectivity | 4.12 | 1,26 | 0.053 | 4.12 | 1,26 | 0.053 |
| **Collateral** | Condition | 6.27 | 1,26 | 0.019 | 6.27 | 1,26 | 0.019 |
| | HP_Head_Strength | 1.28 | 1,26 | 0.269 | 1.28 | 1,26 | 0.269 |
| | CueOnlyForgetting | — | — | — | 10.54 | 1,25 | 0.003 |
| | Condition × Collateral | 0.10 | 1,26 | 0.750 | 0.10 | 1,26 | 0.750 |

*Simple slopes (emtrends) for the Selectivity × Condition interaction: TIS+Cue, b = −18.74, SE = 10.30, t(41.7) = −1.83, p = .075; TIS+Cue delayed, b = 3.03, SE = 10.30, t(41.7) = 0.30, p = .769. HP-Head = left hippocampal head; SE = standard error; df = degrees of freedom*

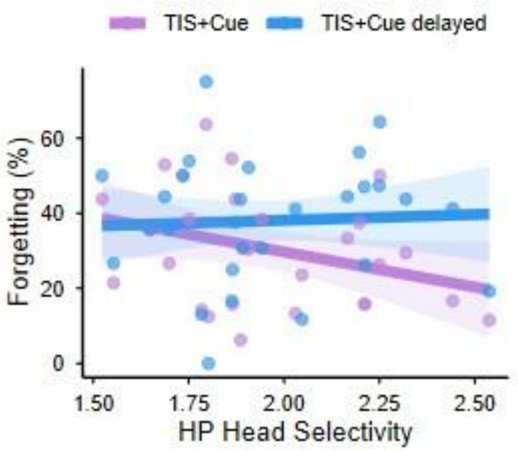


**Fig. S5 | Relationship between TIS selectivity in the hippocampal head and associative memorry.**
Associative memory forgetting showed a trend-level interaction with HP-Head field selectivity and condition (F(1,26) = 4.12, p = 0.053; N=28). Simple slopes: TIS+Cue, b = −18.74, SE = 10.30, t(41.7) = −1.83, p = 0.075; TIS+Cue delayed, b = 3.03, SE = 10.30, t(41.7) = 0.30, p = 0.769 (see Supplementary Table 8). TIS+Cue shown in purple, TIS+Cue delayed in blue; points show individual participants, lines show model-fitted slopes with 95% CI

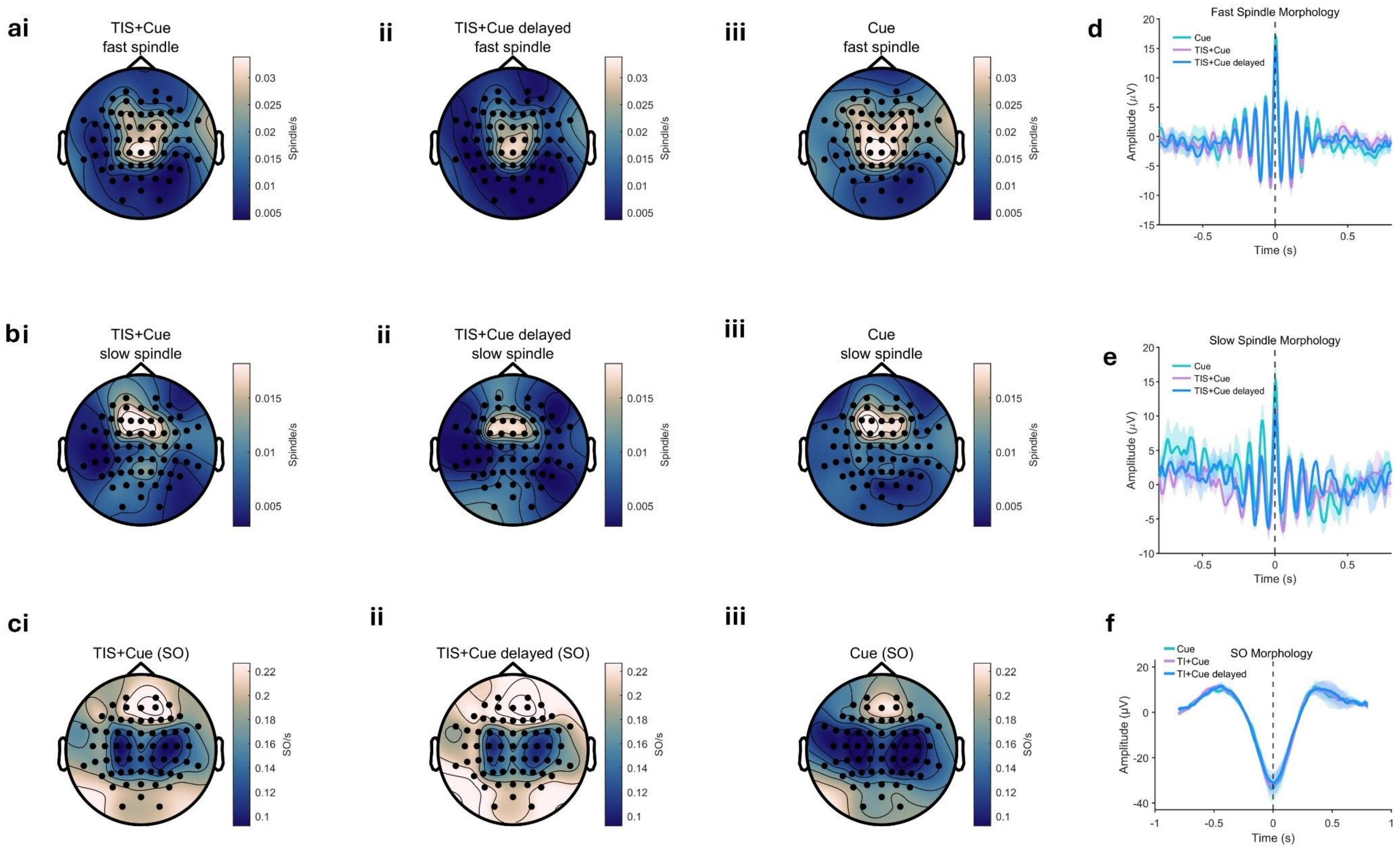


**Fig. S6 | Spindle and slow oscillation distribution and morphology waveforms.**

Topographical distribution of fast spindle density (spindles/s) for TIS+Cue **(i),** TIS+Cue delayed **(ii),** and Cue **(iii). (b)** Topographical distribution of slow spindle density (spindles/s) for TIS+Cue **(i),** TIS+Cue delayed **(ii),** and Cue **(iii). (c)** Topographical distribution of SO density (SO/s) for TIS+Cue **(i),** TIS+Cue delayed **(ii),** and Cue **(iii).** Black dots indicate all 64 electrodes in all topographical plots. **(d)** Average fast spindle morphology across conditions (Cue, TIS+Cue, TIS+Cue delayed), time-locked to spindle peak (t = 0). **(e)** Average slow spindle morphology waveforms across conditions, time-locked to spindle peak (t = 0). **(f)** Average SO morphology waveform across conditions, time-locked to SO negative peak (t = 0). Shaded areas represent ±SEM.

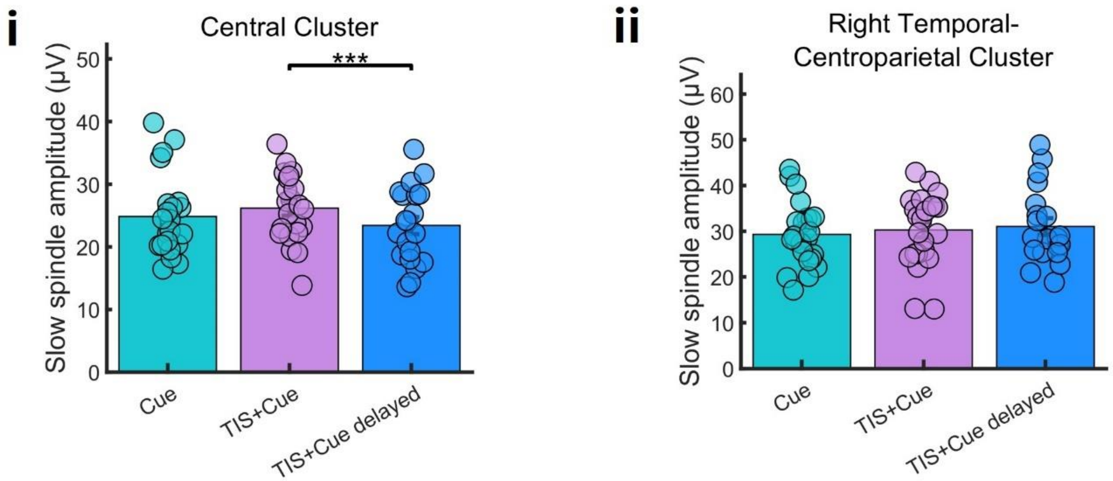


**Fig. | S7 Slow spindle amplitude across conditions.**

Mean slow spindle amplitude (µV) per condition for the central cluster **(i)** and right temporal-centroparietal cluster **(ii).** Post-hoc pairwise comparisons revealed that Cue produced significantly greater slow spindle amplitude than TIS+Cue delayed in the central cluster (**p < .001), whilst no significant differences observed in the right temporal-centroparietal cluster (all p > .05). Dots represent individual participants; bars show mean ± SEM; N=28.

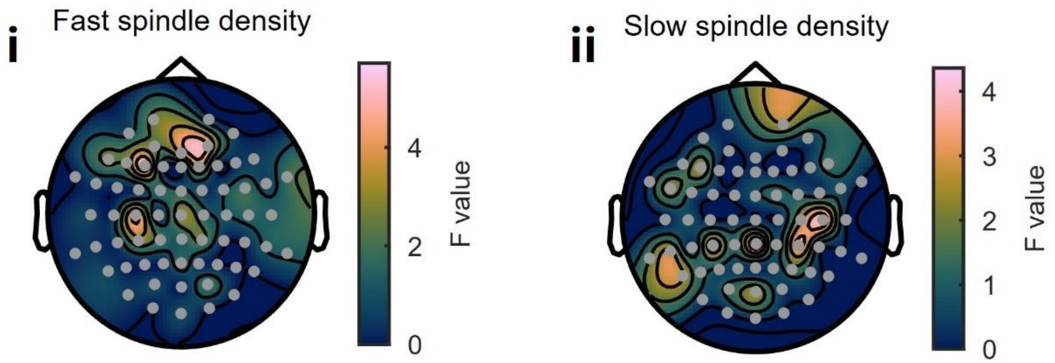


**Fig. | S8 Fast and slow spindle density across conditions.**

Topographical distribution of F-statistics from channel-wise Gamma GLMEs comparing fast spindle density **(i)** and slow spindle density **(ii)** across conditions (Cue, TIS+Cue, TIS+Cue delayed; N=28). No clusters survived cluster-based permutation correction for either measure.

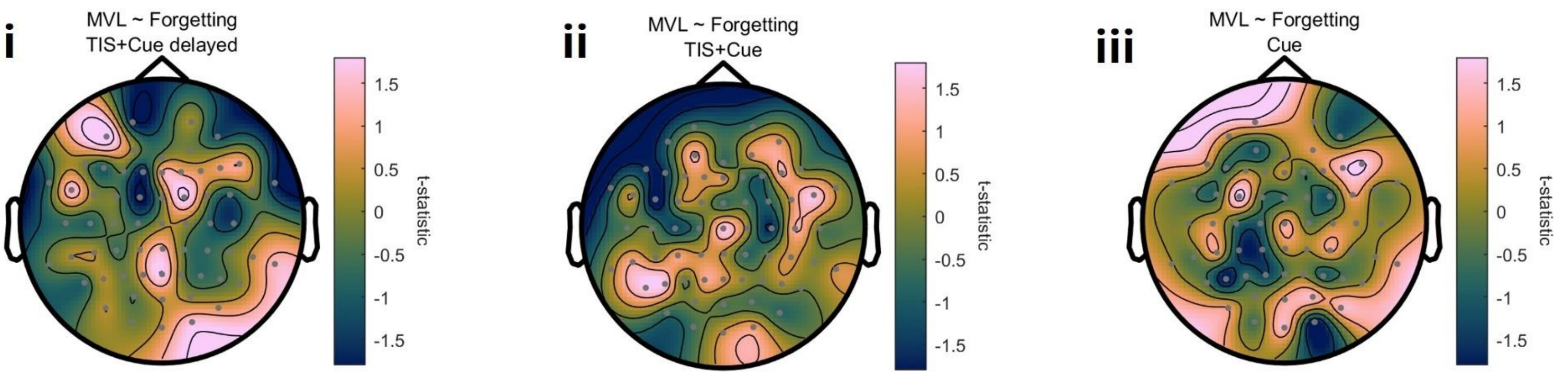


**Fig. | S9 Scalp topography of the relationship between mean vector length (MVL) and associative memory (recall) forgetting.**
Topographical distributions of t-statistics from channel-wise Pearson correlations between SO-spindle coupling strength (MVL) and associative recall forgetting for TIS+Cue **(i),** TIS+Cue delayed **(ii),** and Cue **(iii).** No clusters survived cluster-based permutation correction for any condition.

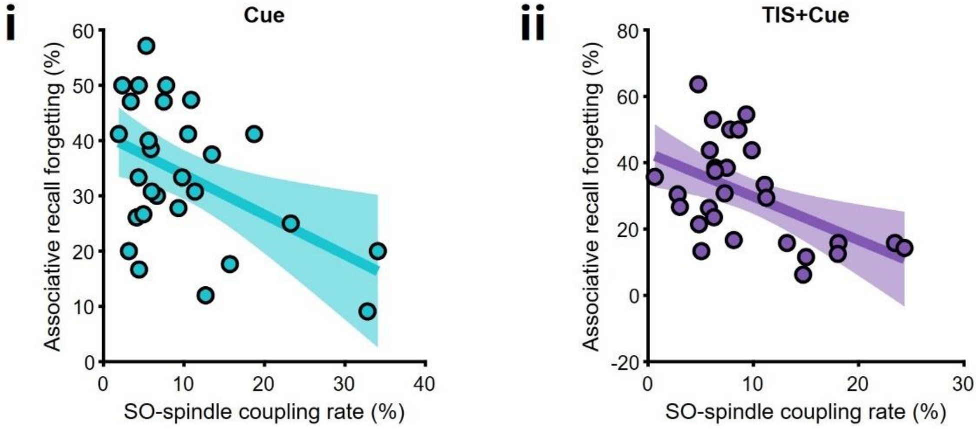


**Fig. S10 | Slow oscillation-spindle coupling rate predicts associative recall forgetting during reactivation.**
SO-spindle coupling rate (averaged over significant channels surviving cluster correction) was associated with reduced forgetting in both Cue ($r = -0.474$, $p = 0.0108$) and TIS+Cue ($r = -0.508$, $p = 0.0057$), ($n = 28$). Dots represent individual participants; lines show fitted linear relationships with shaded bands indicating 95% confidence intervals, N=28.

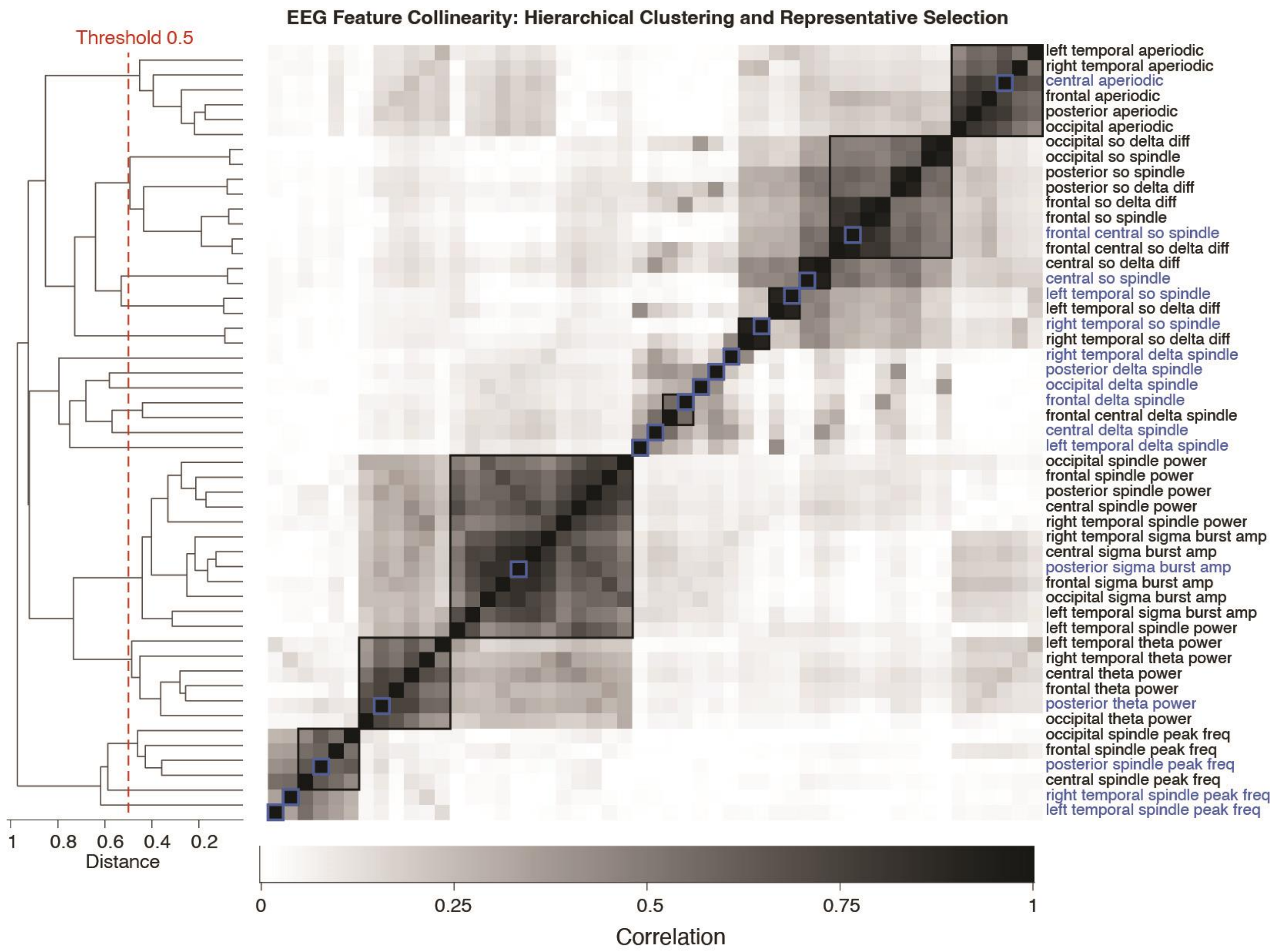


**Fig. S11 | Hierarchical clustering for reducing feature collinearity**.

The features (names shown on the right) are hierarchically clustered (dendrograms on the left) based on their correlation distances. The cell color indicates the strength of correlation between features, with darker cells indicating stronger correlations. The features are clustered at a threshold of 0.5 (red dashed line in dendrogram), yielding 16 clusters. The features within each cluster are enclosed by a large black square, with cluster centroid features highlighted in blue squares.

**Table S7 | Path A coefficients: LME post-hoc pairwise contrasts for oscillatory features across scalp regions.**

| Region | Feature | Contrast | β | SE | t | df | p | p (FDR) |
|---|---|---|---|---|---|---|---|---|
| **Frontal** | Delta → spindle PAC | TIS+Cue vs Cue | 0.0225 | 0.0354 | 0.636 | 3197 | 0.525 | 0.84 |
| **Central** | SO → spindle PAC | TIS+Cue vs Cue | 0.056 | 0.035 | 1.6 | 3182.4 | 0.11 | 0.351 |
| **Central** | Delta → spindle PAC | TIS+Cue vs Cue | -0.0647 | 0.0353 | -1.829 | 3196.5 | 0.0675 | 0.275 |
| **Central** | Spectral slope | TIS+Cue vs Cue | **0.1197** | 0.0345 | 3.473 | 3175.9 | **5.23e-04** | **0.0084** |
| **L Temporal** | Spindle peak frequency | TIS+Cue vs Cue | -0.0021 | 0.0345 | -0.061 | 3176.1 | 0.951 | 0.965 |
| **L Temporal** | SO → spindle PAC | TIS+Cue vs Cue | 0.0644 | 0.0354 | 1.821 | 3196.7 | 0.0687 | 0.275 |
| **L Temporal** | Delta → spindle PAC | TIS+Cue vs Cue | -0.0016 | 0.0354 | -0.044 | 3196.6 | 0.965 | 0.965 |
| **R Temporal** | Spindle peak frequency | TIS+Cue vs Cue | -0.009 | 0.0345 | -0.261 | 3175.7 | 0.794 | 0.965 |
| **R Temporal** | SO → spindle PAC | TIS+Cue vs Cue | 0.034 | 0.0353 | 0.962 | 3192.1 | 0.336 | 0.661 |
| **R Temporal** | Delta → spindle PAC | TIS+Cue vs Cue | -0.0316 | 0.0354 | -0.893 | 3197 | 0.372 | 0.661 |
| **Posterior** | Spindle peak frequency | TIS+Cue vs Cue | **-0.0955** | 0.0334 | -2.859 | 3173.4 | **0.0043** | **0.0342** |
| **Posterior** | Theta power | TIS+Cue vs Cue | 0.006 | 0.0343 | 0.174 | 3176.7 | 0.862 | 0.965 |
| **Posterior** | Sigma burst amplitude | TIS+Cue vs Cue | 0.0492 | 0.0347 | 1.417 | 3180 | 0.156 | 0.417 |
| **Posterior** | Delta → spindle PAC | TIS+Cue vs Cue | 0.0403 | 0.0354 | 1.14 | 3196.6 | 0.254 | 0.581 |
| **Occipital** | Delta → spindle PAC | TIS+Cue vs Cue | -0.0104 | 0.0354 | -0.294 | 3196.1 | 0.769 | 0.965 |
| **Frontal → Central** | SO → spindle PAC | TIS+Cue vs Cue | 0.0057 | 0.0348 | 0.164 | 3180.1 | 0.869 | 0.965 |
| **Frontal** | Delta → spindle PAC | TIS+Cue Delayed vs Cue | 0.0072 | 0.0359 | 0.201 | 3095 | 0.84 | 0.929 |
| **Central** | SO → spindle PAC | TIS+Cue Delayed vs Cue | -0.0031 | 0.0354 | -0.089 | 3078.8 | 0.929 | 0.929 |
| **Central** | Delta → spindle PAC | TIS+Cue Delayed vs Cue | -0.0316 | 0.0359 | -0.88 | 3094.9 | 0.379 | 0.866 |
| **Central** | Spectral slope | TIS+Cue Delayed vs Cue | **0.2022** | 0.0345 | 5.856 | 3073.4 | **5.26e-09** | **8.41e-08** |
| **L Temporal** | Spindle peak frequency | TIS+Cue Delayed vs Cue | 0.0347 | 0.0349 | 0.994 | 3073.7 | 0.321 | 0.855 |
| **L Temporal** | SO → spindle PAC | TIS+Cue Delayed vs Cue | 0.0514 | 0.0359 | 1.431 | 3094.5 | 0.153 | 0.656 |
| **L Temporal** | Delta → spindle PAC | TIS+Cue Delayed vs Cue | -0.0138 | 0.0359 | -0.384 | 3095 | 0.701 | 0.929 |
| **R Temporal** | Spindle peak frequency | TIS+Cue Delayed vs Cue | 0.0097 | 0.0349 | 0.277 | 3073.5 | 0.782 | 0.929 |
| **R Temporal** | SO → spindle PAC | TIS+Cue Delayed vs Cue | -0.0055 | 0.0358 | -0.155 | 3086.9 | 0.877 | 0.929 |
| **R Temporal** | Delta → spindle PAC | TIS+Cue Delayed vs Cue | -0.0272 | 0.0359 | -0.756 | 3095 | 0.449 | 0.899 |
| **Posterior** | Spindle peak frequency | TIS+Cue Delayed vs Cue | -0.0415 | 0.0343 | -1.21 | 3071.2 | 0.227 | 0.725 |
| **Posterior** | Theta power | TIS+Cue Delayed vs Cue | 0.0665 | 0.035 | 1.899 | 3074.5 | 0.0577 | 0.462 |
| **Posterior** | Sigma burst amplitude | TIS+Cue Delayed vs Cue | 0.0214 | 0.0352 | 0.609 | 3077.8 | 0.543 | 0.929 |
| **Posterior** | Delta → spindle PAC | TIS+Cue Delayed vs Cue | 0.019 | 0.0359 | 0.528 | 3095 | 0.597 | 0.929 |
| **Occipital** | Delta → spindle PAC | TIS+Cue Delayed vs Cue | 0.0144 | 0.0359 | 0.399 | 3095 | 0.69 | 0.929 |
| **Frontal → Central** | SO → spindle PAC | TIS+Cue Delayed vs Cue | -0.049 | 0.0352 | -1.392 | 3077.2 | 0.164 | 0.656 |
| **Frontal** | Delta → spindle PAC | TIS+Cue vs TIS+Cue Delayed | 0.0154 | 0.0357 | 0.433 | 3141.8 | 0.665 | 0.793 |
| **Central** | SO → spindle PAC | TIS+Cue vs TIS+Cue Delayed | 0.0568 | 0.0354 | 1.607 | 3128.4 | 0.108 | 0.433 |
| **Central** | Delta → spindle PAC | TIS+Cue vs TIS+Cue Delayed | -0.0302 | 0.0357 | -0.847 | 3146 | 0.397 | 0.747 |
| **Central** | Spectral slope | TIS+Cue vs TIS+Cue Delayed | **-0.0826** | 0.0345 | -2.393 | 3122.6 | **0.0168** | 0.269 |
| **L Temporal** | Spindle peak frequency | TIS+Cue vs TIS+Cue Delayed | -0.0389 | 0.0347 | -1.121 | 3123 | 0.263 | 0.655 |
| **L Temporal** | SO → spindle PAC | TIS+Cue vs TIS+Cue Delayed | 0.0117 | 0.0357 | 0.327 | 3141.1 | 0.744 | 0.793 |
| **L Temporal** | Delta → spindle PAC | TIS+Cue vs TIS+Cue Delayed | 0.012 | 0.0357 | 0.338 | 3146 | 0.736 | 0.793 |
| **R Temporal** | Spindle peak frequency | TIS+Cue vs TIS+Cue Delayed | -0.0202 | 0.0346 | -0.583 | 3122.1 | 0.56 | 0.747 |
| **R Temporal** | SO → spindle PAC | TIS+Cue vs TIS+Cue Delayed | 0.038 | 0.0356 | 1.066 | 3135.7 | 0.287 | 0.655 |
| **R Temporal** | Delta → spindle PAC | TIS+Cue vs TIS+Cue Delayed | -0.0045 | 0.0357 | -0.125 | 3138.2 | 0.9 | 0.9 |
| **Posterior** | Spindle peak frequency | TIS+Cue vs TIS+Cue Delayed | -0.0624 | 0.0337 | -1.855 | 3119.8 | 0.0637 | 0.402 |
| **Posterior** | Theta power | TIS+Cue vs TIS+Cue Delayed | -0.061 | 0.0343 | -1.778 | 3120 | 0.0754 | 0.402 |
| **Posterior** | Sigma burst amplitude | TIS+Cue vs TIS+Cue Delayed | 0.0258 | 0.0345 | 0.746 | 3123.4 | 0.456 | 0.747 |

| Region | Feature | Contrast | β | SE | t | df | p | p (FDR) |
|---|---|---|---|---|---|---|---|---|
| **Posterior** | Delta → spindle PAC | TIS+Cue vs TIS+Cue Delayed | 0.0215 | 0.0357 | 0.602 | 3146 | 0.547 | 0.747 |
| **Occipital** | Delta → spindle PAC | TIS+Cue vs TIS+Cue Delayed | -0.0241 | 0.0357 | -0.675 | 3142.8 | 0.5 | 0.747 |
| **Frontal → Central** | SO → spindle PAC | TIS+Cue vs TIS+Cue Delayed | 0.0444 | 0.035 | 1.269 | 3125.4 | 0.204 | 0.654 |

*β = standardised coefficient; SE = standard error; df = degrees of freedom; p = uncorrected p-value; p (FDR) = Benjamini–Hochberg corrected. Bold = p < 0.05. PAC = phase–amplitude coupling; SO = slow oscillation.*

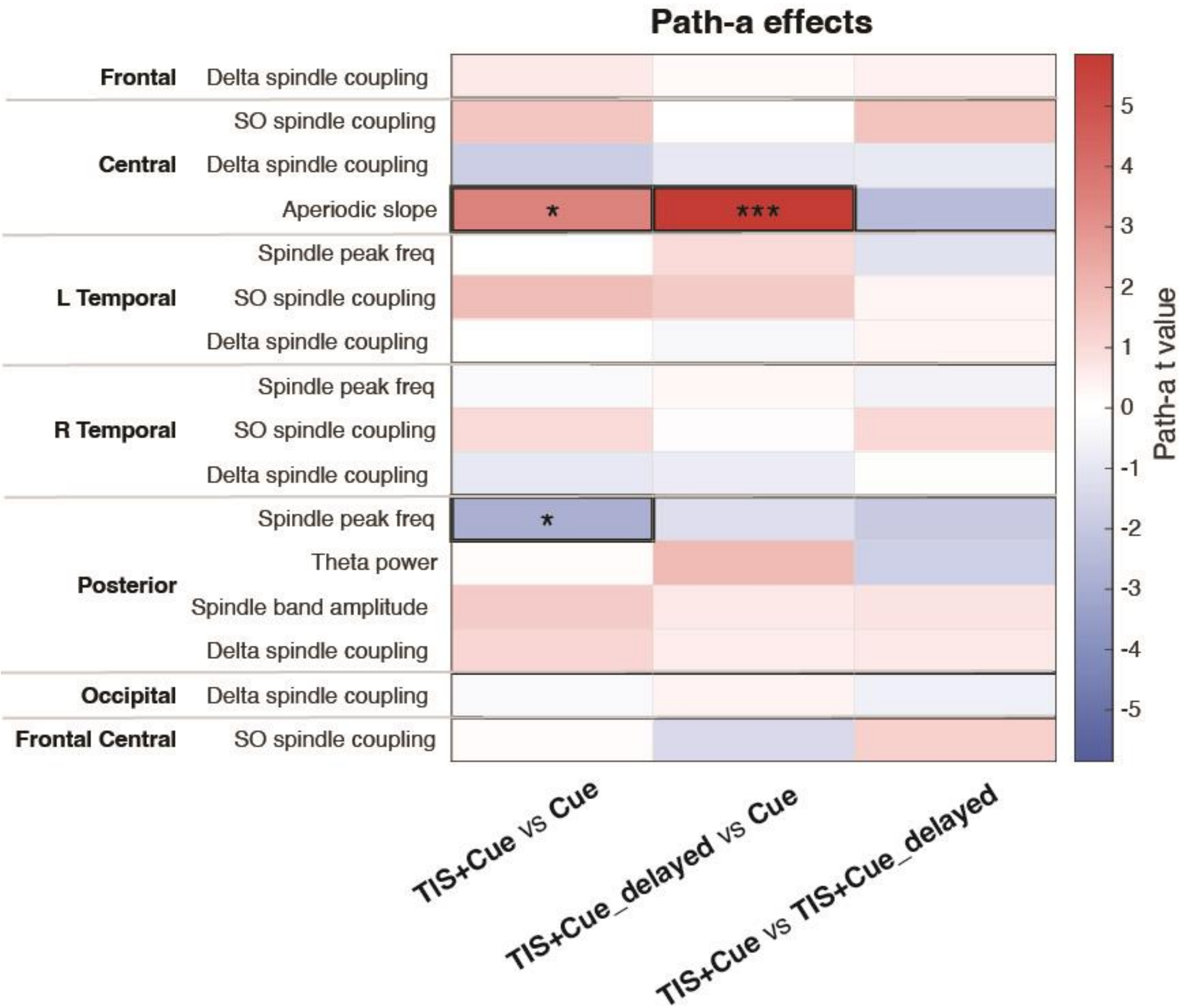


**Fig. S12 | Treatment effects on EEG features (path-a)**

Path-a t-values (effect of condition on each feature) shown as a feature × contrast heatmap. Features (names on the left) are grouped by region; the three columns are the pairwise contrasts (TIS+Cue vs Cue, TIS+Cue_Delayed vs Cue, TIS+Cue vs TIS+Cue_Delayed). Cell colour shows the path-a t-value (red = positive, blue = negative). Asterisks and black outlines mark significance (* $p < 0.05$, *** $p < 0.001$, FDR corrected).

**Table S8 | Mediation analysis: oscillatory features mediating TIS effects on associative memory.**

| Region | Feature | Contrast | N | Path-b β | Path-b p | Path-b p (FDR) | ACME | ACME SD | Norm. ACME | QB p | QB p (FDR) | Perm. p | ADE | Prop. med. | Joint sing. |
|---|---|---|---|---|---|---|---|---|---|---|---|---|---|---|---|
| Frontal | Delta → spindle PAC | TIS+Cue vs Cue | 3312 | 0.0249 | 0.573 | 0.655 | 2.70e-05 | 1.36e-04 | 0.1979 | 0.856 | 0.98 | 0.318 | 0.0228 | 3.00e-04 | False |
| Central | SO → spindle PAC | TIS+Cue vs Cue | 3312 | -0.056 | 0.323 | 0.47 | -1.03e-04 | 3.37e-04 | -0.3047 | 0.77 | 0.98 | 0.941 | 0.0226 | -0.0013 | False |
| Central | Delta → spindle PAC | TIS+Cue vs Cue | 3312 | -0.0598 | 0.225 | 0.359 | 2.33e-04 | 3.04e-04 | 0.7682 | 0.374 | 0.98 | **0.044** | 0.0229 | 0.0061 | False |
| Central | Spectral slope | TIS+Cue vs Cue | 3312 | -0.003 | 0.942 | 0.996 | -1.50e-05 | 2.84e-04 | -0.0524 | 0.96 | 0.98 | 0.744 | 0.0232 | 0 | False |
| L Temporal | Spindle peak frequency | TIS+Cue vs Cue | 3312 | **0.1006** | **0.0181** | 0.145 | 2.17e-04 | 5.90e-04 | 0.3671 | 0.71 | 0.98 | 0.496 | 0.023 | 0.0071 | False |
| L Temporal | SO → spindle PAC | TIS+Cue vs Cue | 3312 | 0.0898 | 0.0508 | 0.271 | 3.64e-04 | 3.45e-04 | 1.0564 | 0.206 | 0.98 | **0.03** | 0.0222 | 0.0115 | False |
| L Temporal | Delta-spindle PAC | TIS+Cue vs Cue | 3312 | 0.0612 | 0.161 | 0.322 | -1.00e-06 | 2.01e-04 | -0.0057 | 0.96 | 0.98 | 0.474 | 0.0232 | -1.00e-04 | False |
| R Temporal | Spindle peak frequency | TIS+Cue vs Cue | 3312 | -2.00e-04 | 0.996 | 0.996 | -2.00e-06 | 2.37e-04 | -0.0096 | 0.98 | 0.98 | 0.583 | 0.0226 | 2.00e-04 | False |
| R Temporal | SO → spindle PAC | TIS+Cue vs Cue | 3312 | 0.0727 | 0.101 | 0.313 | 1.18e-04 | 2.75e-04 | 0.427 | 0.662 | 0.98 | 0.169 | 0.0238 | 0.0024 | False |
| R Temporal | Delta → spindle PAC | TIS+Cue vs Cue | 3312 | 0.0331 | 0.437 | 0.582 | -6.40e-05 | 1.74e-04 | -0.3666 | 0.722 | 0.98 | 0.828 | 0.0231 | -0.001 | False |
| Posterior | Spindle peak frequency | TIS+Cue vs Cue | 3312 | -0.0587 | 0.218 | 0.359 | 1.49e-04 | 5.38e-04 | 0.2764 | 0.798 | 0.98 | **0.013** | 0.0221 | 0.0024 | False |
| Posterior | Theta power | TIS+Cue vs Cue | 3312 | 0.0638 | 0.161 | 0.322 | 4.70e-05 | 4.33e-04 | 0.1077 | 0.936 | 0.98 | 0.406 | 0.0232 | 4.00e-04 | False |
| Posterior | Sigma burst amplitude | TIS+Cue vs Cue | 3312 | **0.127** | **0.0024** | **0.0384** | 2.99e-04 | 6.32e-04 | 0.4726 | 0.612 | 0.98 | 0.11 | 0.0225 | 0.0101 | False |
| Posterior | Delta → spindle PAC | TIS+Cue vs Cue | 3312 | 0.074 | 0.117 | 0.313 | 1.86e-04 | 2.66e-04 | 0.6999 | 0.462 | 0.98 | 0.128 | 0.0226 | 0.0049 | False |
| Occipital | Delta → spindle PAC | TIS+Cue vs Cue | 3312 | -0.0715 | 0.107 | 0.313 | 3.60e-05 | 2.37e-04 | 0.1512 | 0.858 | 0.98 | 0.331 | 0.0229 | 6.00e-04 | False |
| Frontal → Central | SO → spindle PAC | TIS+Cue vs Cue | 3312 | -0.032 | 0.505 | 0.622 | 2.40e-05 | 2.69e-04 | 0.0884 | 0.922 | 0.98 | 0.65 | 0.0233 | 3.00e-04 | False |
| Frontal | Delta → spindle PAC | TIS+Cue Delayed vs Cue | 3218 | -0.0571 | 0.193 | 0.441 | -2.70e-05 | 1.97e-04 | -0.1387 | 0.87 | 0.934 | 0.544 | -0.055 | 2.00e-04 | False |
| Central | SO → spindle PAC | TIS+Cue Delayed vs Cue | 3218 | 0.0435 | 0.439 | 0.722 | -5.90e-05 | 3.25e-04 | -0.18 | 0.846 | 0.934 | 0.617 | -0.0557 | 5.00e-04 | False |
| Central | Delta → spindle PAC | TIS+Cue Delayed vs Cue | 3218 | -0.0257 | 0.595 | 0.722 | 4.00e-05 | 1.72e-04 | 0.2291 | 0.854 | 0.934 | 0.163 | -0.0551 | -3.00e-04 | False |
| Central | Spectral slope | TIS+Cue Delayed vs Cue | 3218 | 0.028 | 0.506 | 0.722 | 1.87e-04 | 4.25e-04 | 0.4415 | 0.688 | 0.934 | **9.99e-04** | -0.0548 | -0.0018 | False |
| L Temporal | Spindle peak frequency | TIS+Cue Delayed vs Cue | 3218 | **0.13** | **0.0023** | **0.0374** | 4.70e-04 | 7.90e-04 | 0.5951 | 0.522 | 0.934 | 0.192 | -0.0541 | -0.0081 | False |
| L Temporal | SO → spindle PAC | TIS+Cue Delayed vs Cue | 3218 | 0.0088 | 0.845 | 0.845 | 2.80e-05 | 1.78e-04 | 0.1586 | 0.866 | 0.934 | **0.032** | -0.0541 | -2.00e-04 | False |
| L Temporal | Delta → spindle PAC | TIS+Cue Delayed vs Cue | 3218 | -0.0209 | 0.632 | 0.722 | 1.00e-05 | 1.42e-04 | 0.0727 | 0.926 | 0.934 | 0.397 | -0.0541 | -1.00e-04 | False |

| Region | Feature | Contrast | N | Path-b β | Path-b p | Path-b p (FDR) | ACME | ACME SD | Norm. ACME | QB p | QB p (FDR) | Perm. p | ADE | Prop. med. | Joint sing. |
|---|---|---|---|---|---|---|---|---|---|---|---|---|---|---|---|
| R Temporal | Spindle peak frequency | TIS+Cue Delayed vs Cue | 3218 | -0.0742 | 0.0809 | 0.259 | -2.48e-04 | 5.47e-04 | -0.4523 | 0.64 | 0.934 | 0.512 | -0.0549 | 0.0032 | False |
| R Temporal | SO → spindle PAC | TIS+Cue Delayed vs Cue | 3218 | 0.0509 | 0.246 | 0.493 | -4.20e-05 | 2.62e-04 | -0.1597 | 0.866 | 0.934 | 0.541 | -0.0541 | 3.00e-04 | False |
| R Temporal | Delta → spindle PAC | TIS+Cue Delayed vs Cue | 3218 | 0.0776 | 0.0664 | 0.259 | -1.20e-04 | 2.54e-04 | -0.473 | 0.634 | 0.934 | 0.769 | -0.0552 | 0.0016 | False |
| Posterior | Spindle peak frequency | TIS+Cue Delayed vs Cue | 3218 | -0.0139 | 0.767 | 0.818 | 3.50e-05 | 3.55e-04 | 0.0971 | 0.934 | 0.934 | 0.0719 | -0.055 | -1.00e-04 | False |
| Posterior | Theta power | TIS+Cue Delayed vs Cue | 3218 | 0.0339 | 0.46 | 0.722 | 1.80e-04 | 4.05e-04 | 0.4454 | 0.66 | 0.934 | **0.041** | -0.0554 | -0.0016 | False |
| Posterior | Sigma burst amplitude | TIS+Cue Delayed vs Cue | 3218 | **0.1101** | **0.0094** | 0.0751 | 2.70e-05 | 5.92e-04 | 0.0457 | 0.926 | 0.934 | 0.251 | -0.0546 | -9.00e-04 | False |
| Posterior | Delta → spindle PAC | TIS+Cue Delayed vs Cue | 3218 | **0.1074** | **0.0249** | 0.133 | 1.32e-04 | 3.46e-04 | 0.3818 | 0.686 | 0.934 | 0.383 | -0.0561 | -0.0018 | False |
| Occipital | Delta → spindle PAC | TIS+Cue Delayed vs Cue | 3218 | -0.0257 | 0.566 | 0.722 | -2.20e-05 | 1.37e-04 | -0.1604 | 0.86 | 0.934 | 0.571 | -0.0552 | 2.00e-04 | False |
| Frontal → Central | SO → spindle PAC | TIS+Cue Delayed vs Cue | 3218 | -0.0794 | 0.0977 | 0.26 | 3.40e-04 | 5.33e-04 | 0.638 | 0.474 | 0.934 | 0.0729 | -0.0541 | -0.0045 | False |
| Frontal | Delta → spindle PAC | TIS+Cue vs TIS+Cue Delayed | 3274 | -0.0235 | 0.591 | 0.945 | -1.70e-05 | 1.43e-04 | -0.1214 | 0.878 | 0.994 | 0.595 | 0.0833 | -1.00e-04 | False |
| Central | SO → spindle PAC | TIS+Cue vs TIS+Cue Delayed | 3274 | 0.0242 | 0.676 | 0.945 | 6.60e-05 | 3.20e-04 | 0.2069 | 0.836 | 0.994 | 0.125 | 0.0824 | 4.00e-04 | False |
| Central | Delta → spindle PAC | TIS+Cue vs TIS+Cue Delayed | 3274 | 0.0215 | 0.664 | 0.945 | -4.30e-05 | 1.68e-04 | -0.253 | 0.794 | 0.994 | 0.899 | 0.0815 | -2.00e-04 | False |
| Central | Spectral slope | TIS+Cue vs TIS+Cue Delayed | 3274 | -0.0039 | 0.927 | 0.945 | 1.90e-05 | 4.17e-04 | 0.0457 | 0.994 | 0.994 | 0.63 | 0.0823 | 0 | False |
| L Temporal | Spindle peak frequency | TIS+Cue vs TIS+Cue Delayed | 3274 | **0.0937** | **0.0276** | 0.181 | 5.00e-06 | 5.31e-04 | 0.0091 | 0.978 | 0.994 | 0.872 | 0.0816 | -1.00e-04 | False |
| L Temporal | SO → spindle PAC | TIS+Cue vs TIS+Cue Delayed | 3274 | -0.0387 | 0.401 | 0.917 | -2.40e-05 | 1.76e-04 | -0.1337 | 0.91 | 0.994 | 0.659 | 0.0824 | -1.00e-04 | False |
| L Temporal | Delta → spindle PAC | TIS+Cue vs TIS+Cue Delayed | 3274 | -0.0373 | 0.396 | 0.917 | -2.80e-05 | 1.63e-04 | -0.1704 | 0.882 | 0.994 | 0.638 | 0.0821 | -1.00e-04 | False |
| R Temporal | Spindle peak frequency | TIS+Cue vs TIS+Cue Delayed | 3274 | **-0.0904** | **0.0339** | 0.181 | -1.94e-04 | 5.77e-04 | -0.3363 | 0.732 | 0.994 | 0.321 | 0.0821 | -0.0015 | False |
| R Temporal | SO → spindle PAC | TIS+Cue vs TIS+Cue Delayed | 3274 | 0.0129 | 0.772 | 0.945 | 3.70e-05 | 1.52e-04 | 0.2427 | 0.844 | 0.994 | 0.0959 | 0.0824 | 1.00e-04 | False |
| R Temporal | Delta → spindle PAC | TIS+Cue vs TIS+Cue Delayed | 3274 | -0.0039 | 0.928 | 0.945 | -1.00e-06 | 1.27e-04 | -0.0049 | 0.978 | 0.994 | 0.513 | 0.0827 | 0 | False |

| Region | Feature | Contrast | N | Path-b β | Path-b p | Path-b p (FDR) | ACME | ACME SD | Norm. ACME | QB p | QB p (FDR) | Perm. p | ADE | Prop. med. | Joint sing. |
|---|---|---|---|---|---|---|---|---|---|---|---|---|---|---|---|
| Posterior | Spindle peak frequency | TIS+Cue vs TIS+Cue Delayed | 3274 | 0.0087 | 0.855 | 0.945 | -1.60e-05 | 4.01e-04 | -0.0408 | 0.94 | 0.994 | 0.76 | 0.0822 | -1.00e-04 | False |
| Posterior | Theta power | TIS+Cue vs TIS+Cue Delayed | 3274 | 0.0032 | 0.945 | 0.945 | 1.00e-06 | 3.51e-04 | 0.0021 | 0.994 | 0.994 | 0.432 | 0.0822 | 0 | False |
| Posterior | Sigma burst amplitude | TIS+Cue vs TIS+Cue Delayed | 3274 | 0.0054 | 0.9 | 0.945 | 2.00e-06 | 2.60e-04 | 0.0069 | 0.978 | 0.994 | 0.346 | 0.083 | 0 | False |
| Posterior | Delta → spindle PAC | TIS+Cue vs TIS+Cue Delayed | 3274 | **0.1052** | **0.0265** | 0.181 | 1.47e-04 | 3.23e-04 | 0.4541 | 0.618 | 0.994 | 0.263 | 0.0828 | 0.0014 | False |
| Occipital | Delta → spindle PAC | TIS+Cue vs TIS+Cue Delayed | 3274 | -0.0508 | 0.251 | 0.803 | 7.10e-05 | 2.04e-04 | 0.3465 | 0.702 | 0.994 | 0.248 | 0.0826 | 4.00e-04 | False |
| Frontal → Central | SO → spindle PAC | TIS+Cue vs TIS+Cue Delayed | 3274 | -0.0754 | 0.127 | 0.507 | -1.08e-04 | 4.07e-04 | -0.266 | 0.782 | 0.994 | 0.857 | 0.0825 | -7.00e-04 | False |

*β = standardised coefficient; ACME = average causal mediation effect; Norm. ACME = ACME normalised by total effect; QB = Quasi-Bayesian; Perm. p = permutation-based p-value; ADE = average direct effect; Prop. med. = proportion of total effect mediated; Joint sing. = joint singularity flag. Bold = p < 0.05.*

##### Table S9 | Perceptual sensations and thresholds across participants for conventional tACS and temporal interference stimulation

Individual perceptual thresholds and reported sensations for each participant (ID) during conventional tACS, (5 Hz) and TIS. Prior to the nap, each participant underwent threshold testing starting with tACS to familiarize participants with potential perceptual effects of electrical stimulation. Current intensity was gradually increased in 0.1 mA increments until they reported feeling a sensation. Threshold determination proceeded sequentially across electrode pairs, first e1-e2, then e3-e4.

| | tACS (5 Hz) | | | | TIS CF = 2 and 2.090 kHz, f = 90 Hz | | | |
|---|---|---|---|---|---|---|---|---|
| | e1 – e2 | | e3 – e4 | | e1 – e2 | | e3 – e4 | |
| **ID** | **Threshold (mA)** | **Sensation** | **Threshold (mA)** | **Sensation** | **Threshold (mA)** | **Sensation** | **Threshold (mA)** | **Sensation** |
| **1** | 0.3 | tingling | 0.5 | tingling | 2.5 | tingling | 3 | tingling |
| **2** | 0.1 | tingling | 0.1 | tingling | 1.5 | warm tingling | 2.5 | tingling |
| **3** | 0.1 | tingling | 0.1 | tingling | 1.5 | sharp tingling | 3 | none |
| **4** | 0.5 | tingling | 0.5 | tingling | 2.5 | tingling | 3 | none |
| **5** | 0.1 | tingling | 0.3 | tingling | 2.5 | tingling | 3 | itching |
| **6** | 0.1 | tingling | 0.1 | pulsing | 2.5 | tingling | 3 | pinpricks |
| **7** | 0.3 | tingling | 0.5 | pulsing | 3 | tingling | 3 | tingling |
| **8** | 0.3 | tingling | 0.3 | tingling | 2.5 | tingling | 2.5 | tingling |
| **9** | 0.3 | tingling | 0.7 | tingling | 2.5 | tingling | 2.5 | itching |

| | tACS (5 Hz) | | | | TIS CF = 2 and 2.090 kHz, f = 90 Hz | | | |
|---|---|---|---|---|---|---|---|---|
| **10** | 0.3 | pinpricks | 0.5 | pinpricks | 2.5 | tingling | 3 | none |
| **11** | 0.3 | tingling | 0.3 | pinpricks | 3 | pressure | 3 | pinpricks |
| **12** | 0.5 | pinpricks | 0.3 | tingling | 2 | tingling | 3 | tingling |
| **13** | 0.3 | tingling | 0.5 | pinpricks | 1.5 | vibration | 2.5 | vibration |
| **14** | 0.3 | burning | 0.3 | tingling | 2 | tingling | 3 | vibration |
| **15** | 0.3 | tingling | 0.5 | tingling | 2 | tingling | 3 | none |
| **16** | 0.3 | tingling | 0.5 | tingling | 2.5 | tingling | 3 | none |
| **17** | 0.5 | tingling | 0.5 | pinpricks | 2 | tingling | 3 | None |
| **18** | 0.7 | pinpricks | 0.6 | tingling | 2.5 | tingling | 2.5 | none |
| **19** | 0.5 | tingling | 0.3 | tingling | 3 | sharp tingling | 3 | none |
| **20** | 0.4 | tingling | 0.7 | tingling | 2.5 | tingling | 3 | tingling |
| **21** | 0.5 | tingling | 0.6 | heat | 2.5 | pinpricks | 3 | pinpricks |
| **22** | 0.3 | burning | 0.3 | pinpricks | 3 | moving pressure | 3 | none |
| **23** | 0.5 | pinpricks | 0.6 | tapping | 2.5 | pinpricks | 3 | none |
| **24** | 0.5 | pinpricks | 0.5 | heat | 3 | pressure | 3 | none |
| **25** | 0.5 | Heat | 0.5 | tingling | 2.5 | tingling | 3 | none |
| **26** | 0.7 | pulsing | 0.5 | tingling | 2.5 | tingling | 3 | none |
| **27** | 0.5 | tingling | 0.7 | pinpricks | 2 | Pinpricks | 3 | none |
| **28** | 0.5 | pinpricks | 0.7 | tingling | 2.5 | pressure | 3 | none |

*tACS = transcranial alternating current stimulation; TIS = temporal interference stimulation; e = electrode*

#### Table S10 | Summary of post-stimulation side effects questionnaire for TIS during sleep

Summary statistics for adverse effects questionnaire[1]. Participants rated each sensation between 1 (very weak) and 5 (severe). Average intensity and range calculated over participants who reported the sensation (i.e., rating > 0).

| | Mean Intensity (SD) | Number of incidents > 0 (out of 28) | Range |
|---|---|---|---|
| **Itching** | 2.25 (1.26) | 4 | 1 - 4 |
| **Tingling** | 1.71 (0.76) | 7 | 1 - 3 |
| **Headache** | 1.50 (0.71) | 2 | 1 - 2 |
| **Scalp pain** | 1(0) | 2 | 1 |
| **Neck pain** | 2(0) | 1 | 2 |
| **Burning** | 1(0) | 1 | 1 |
| **Warmth/Heat** | 2.17 (1.33) | 6 | 1 - 4 |
| **Metallic taste** | 1(0) | 1 | 1 |
| **Nervousness/Anxiety** | 1(0) | 1 | 1 |
| **Discomfort** | 1(0) | 6 | 1 |
| **Dizziness** | 3(0) | 1 | 3 |
| **Visual sensations** | 2(0) | 1 | 2 |
| **Other** | 2(0) | 1 | 2 |

*SD, standard deviation; TIS = temporal interference stimulation.*

##### Table S11 | Changes in subjective sleepiness from pre-nap to post-nap

Pre- and post-nap subjective sleepiness measured using the KSS[2]. Participants (N = 28) completed the KSS immediately before the afternoon nap and immediately upon awakening. Values were compared using a paired-samples t-test. The KSS ranges from 1 (very alert) to 9 (extremely sleepy, fighting sleep); lower scores indicate greater alertness.

| | Pre-nap | | Post-nap | | Pre vs Post | | |
|---|---|---|---|---|---|---|---|
| | Mean | SD | Mean | SD | t | DF | p-value |
| KSS | 5.8 | 1.4 | 4.2 | 1.7 | 3.78 | 27 | <0.001 |

*KSS = Karolinska Sleepiness Scale*

##### Table S12 | Subjective sleep quality and restfulness following temporal interference stimulation during nap

Subjective ratings of sleep quality and restfulness upon awakening from nap. Participants (N = 28) rated their overall sleep quality on a 6-point scale and how rested they felt upon awakening on a 5-point scale. Ratings were collected immediately following the experimental nap during which TIS was delivered during slow-wave sleep.

| | Mean (SD) | Scale |
|---|---|---|
| **Sleep quality** | 4.14 (1.11) | 1 = very poor, 6 = excellent |
| **Restfulness upon awakening** | 3.18 (0.98) | 1 = not at all rested, 5 = very well-rested |

*SD, standard deviation; TIS = temporal interference stimulation.*

#### *Supplementary Materials:*

##### Electrode positioning formulae

Electrode positions were placed based on each participant's head circumference measured in centimetres. The half circumference was first computed as circumference ÷ 2, and electrode positions were then derived as follows: e1 = (half circumference ÷ 2) − 2.5; e3 = (half circumference ÷ 2) + 2.5; e2 = (half circumference × 0.2) − 1; e4 = (half circumference × 0.2) + 1 + (half circumference ÷ 2). For example, for a participant with a head circumference of 58 cm (half circumference = 29 cm), electrode positions were: e1 = 12.0 cm, e3 = 17.0 cm, e2 = 4.8 cm, and e4 = 21.3 cm.

#### References


1. Violante, I. R. *et al.* Non-invasive temporal interference electrical stimulation of the human hippocampus. *Nat. Neurosci.* **26**, 1994–2004 (2023).

2. Akerstedt, T. *Subjective and objective sleepiness in the active individual*. vol. 52 (1990).
